# Constant-pH simulation of the human *β*_2_ adrenergic receptor inactivation

**DOI:** 10.1101/2025.06.26.661669

**Authors:** Federico Ballabio, Riccardo Capelli

## Abstract

Understanding the molecular basis of pH-dependent G protein-coupled receptor (GPCR) signaling is crucial for comprehending physiological regulation and drug design. Here, we investigate the human *β*_2_ adrenergic receptor (*β*_2_AR), a prototypical GPCR whose function is sensitive to pH conditions. Employing extensive constant-pH molecular dynamics simulations, we provide a detailed atomistic characterization of *β*_2_AR inactivation across physiologically relevant pH values (4–9). Our simulations reveal that *β*_2_AR inactivation is closely linked to protonation events at critical residues, notably E268^6×30^ involved in the ionic lock formation. Furthermore, we find that inactivation occurs without direct sodium binding to the ion-binding pocket around residue D79^2×50^. Instead, sodium ions predominantly interact with D113^3×32^, effectively blocking deeper entry toward the traditional binding site. These results challenge existing mechanistic models and highlight the necessity of accurately modeling electrostatics in GPCR simulations. Our findings underscore the potential of constant-pH methodologies to advance the understanding of GPCR dynamics, influencing both fundamental biology and therapeutic strategies.

## Introduction

G protein-coupled receptors (GPCRs) constitute a vast and versatile family of membrane proteins that play a pivotal role in cellular communication.^1^ These receptors detect a wide array of external signals –ranging from hormones and neurotransmitters to sensory stimuli– and convert them into intracellular responses. Their characteristic seven-transmembrane helical structure not only facilitates the transmission of extracellular signals, but also ensures precise activation of downstream signaling cascades, primarily through interactions with heterotrimeric G proteins. Given their central importance in numerous physiological processes, GPCRs have become prominent targets in drug discovery and therapeutic intervention. ^2,3^ In this fundamental family of proteins, one of the first members for which the structure was experimentally determined was the *β*_2_ adrenergic receptor (*β*_2_AR), which is linked to the modulation of cardiovascular and respiratory responses as well as metabolic regulation. Since its structure was first resolved in 2007,^4,5^ this receptor has served as a cornerstone for understanding GPCR architecture and activation mechanisms. Its structural characterization and a series of NMR experiments^6,7^ have paved the way for extensive studies investigating ligand binding, G protein coupling, and subsequent signaling pathways, thereby providing critical insights into GPCR-mediated signal transduction.^8,9^

In recent years, classical molecular dynamics (MD) simulations have emerged as an essential tool in computational structural biology. MD simulations complement experimental techniques by providing an atomistic and dynamic view of protein behavior, thereby elucidating the transient conformational states and molecular interactions that underlie GPCR function. This approach captures the subtle changes in receptor structure that occur during activation/inactivation, offering a dynamic perspective that static experimental structures cannot fully reveal. In this regard, *β*_2_AR made no exception: since its structural determination, it was extensively studied by means of MD simulations. In particular, the spontaneous formation of the ionic lock in the inactive state was observed *in silico* almost immediately after the release of the crystal structure, on a timescale of hundreds of ns. ^10,11^ This was followed by long-timescale simulations that captured the activation and inactivation profiles, highlighting the microswitches involved in the process. ^12^ Experimentally, the activity of *β*_2_AR has been shown to be pH-dependent: at acidic pH (6.5), the receptor results more active, while at slightly more basic pH (8) *β*_2_AR appears mainly inactive.^13^ This effect is usually associated with the different protonation states of key residues and their interaction with sodium. These factors drive the inactivation in multiple receptors belonging to the same family. ^14,15^ The binding process of sodium ions in *β*_2_AR to the D^2×50^ (the superscript refers to Ballesteros-Weinstein numbering,^16^ following the GPCRdb classification^17^) has been extensively studied *in silico*, generally recognizing its key role as driver for conformational changes in general, and activation/inactivation^18^ in particular. In multiple computational works it has been suggested that sodium ions interactions trigger and stabilize the inactivation of *β*_2_AR,^19,20^ and a precise pathway for sodium binding has been identified and characterized.^21^ However, classical MD simulations fix the protonation states of titrable sites. This does not allow for an on-the-fly change in protonation, which has been suggested to be fundamental in regulating the activation/inactivation mechanism.^10,11^

To take into account the possibility of a dynamic protonation state in classical simulations, two main families of approach have been developed: (i) the discrete constant-pH framework, where a hybrid MD/Monte Carlo (MC) approach is devoted in changing the protonation state of titrable residues after a Poisson-Boltzmann environment evaluation (implemented in multiple ways^22–25)^ or (ii) a continuous constant-pH framework, where a variable, usually called *λ*, interpolates between the charged and neutral states of every titrable site, being influenced by the pH value and background conditions (also in this case, with multiple implementations available^26–29)^. Among these options, we choose the GROMACS-based implementation of the continuous constant-pH simulations proposed by Aho and coworkers^29,30^ for its scalability with respect to the number of titrable sites involved; such approach has been recently employed in the study of a proton sensing GPCRs.^31^

Here we performed microsecond-long constant-pH MD simulations starting from the active state of *β*_2_AR under different pH conditions, covering a range compatible with the known experimental findings (from 4 to 9). Surprisingly, our simulations revealed a more nuanced picture than a purely pH-dependent activation. We observed substantial maintenance of the active state at low pHs and a consistent inactivation across different replicas at higher pH. Furthermore, we never observed spontaneous binding of Na^+^ ions to D^2×50^, despite the presence of a clear inactivation process that involved multiple microswitches already identified in previous works.^32^

These findings highlight the need for better modeling of electrostatics effects (protonation and polarization effects) in the simulation of molecular systems of biological interest.

## Methods

### System preparation

The active-state model of *β*_2_AR was generated using the AlphaFold3 web server.^33^ To bias the prediction toward the active conformation, the sequence of the active state-stabilizing Nb80 nanobody (residues 2-122) from the *β*_2_AR–Nb80 complex (PDB ID: 3P0G^34^) was included alongside the receptor sequence (residues D23^N-term^–344^C-term^). The best predicted model (iPTM score = 0.87, PTM score = 0.85) was selected, and the nanobody was subsequently removed. The receptor conformation was then used as the starting structure for our calculations. To validate that the predicted model adopted an active-state conformation, its C*_α_* atoms from transmembrane helices TM1–TM7 were aligned to the corresponding C*_α_*atoms of *β*_2_AR in the nanobody-bound experimental structure 3P0G^34^ (RMSD_TM1-7_ of 0.6 Å) and the G protein-bound experimental structure 3SN6^35^(RMSD_TM1-7_ of 0.9 Å), Figure S1 in the Supporting Information.

### Setup of constant-pH simulations

To prepare our simulation, we took the active state *β*_2_AR system prepared as previously described and, using CHARMM-GUI,^36^ we embedded it in a POPC bilayer and solvated it, having a cubic box of size 89 × 89 × 119 Å. The membrane, the protein, and the ions were parameterized with the CHARMM36m force field,^37^ and the solvent was modeled using the TIP3P water model.^38^

After constructing the all-atom system, we used the phbuilder tool^39^ to prepare the titrable model of *β*_2_AR at different pH values (4, 5, 6, 7, 8, and 9), setting as titrable all the HIS, ARG, LYS, ASP, and GLU residues. We neutralized the box using sodium chloride, fixing its concentration to 0.15 M, also adding 20 buffer particles to maintain the neutrality in case of change in the protonation state of the titrable residues.

### Molecular dynamics simulations

For each of the six pH values, we performed a minimization and equilibration protocol following the CHARMM-GUI pipeline. Namely, we first performed a steepest descent minimization (until we reach a maximum force of 1000 kJ/mol/nm), and a series of equilibration steps with constraints, keeping the *λ*-states of the titrable residues fixed (see table 1).

**Table 1:**
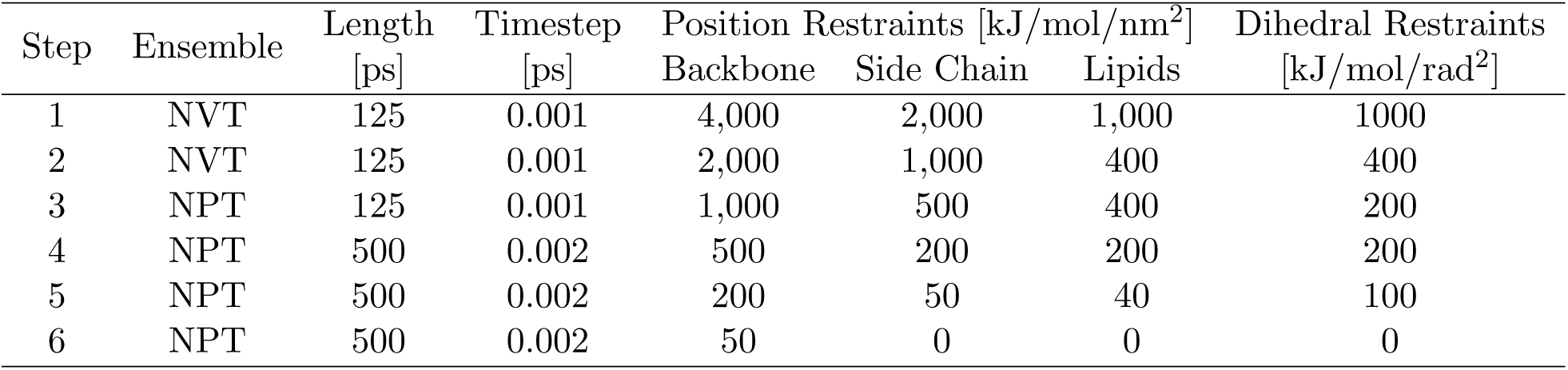
Equilibration protocol.

After the equilibration protocol, we ran five independent replicas, each 1 *µ*s-long, for each pH value, re-initializing the initial velocities. A total of 30 *µ*s of trajectories were collected. In such simulations, the *λ*-parameters that account for the protonation states of titrable residues were allowed to change based on the surrounding environment. For all the simulations, the short-range electrostatics cutoff was set to 1.2 nm, while the long-range was modeled via Particle-Mesh Ewald summation,^40^ in agreement with the prescription of the CHARMM36m force field.^37^ The temperature was kept constant to 310 K via the velocity rescale thermostat^41^ (coupling time of 1 ps), while the pressure (for NPT parts of the protocol) was kept constant to 1 bar using the cell rescale barostat^42^ (coupling time of 5 ps, compressibility of 4.5 · 10*^−^*^5^ bar^-1^). To allow the use of a 2 fs timestep, we applied the LINCS algorithm^43^ to the H-bonds. All the simulations were performed using GROMACS 2021^44^ in the constant-pH-modified version^29,30^ (retrieved at https://gitlab.com/gromacs-constantph/constantph).

### Microswitches definitions

To evaluate the activation state of the receptor during the simulation, we considered four molecular microswitches defined in previous works: ^6,12,45^ (i) the ionic lock distance (distance between R131^3×50^ and L272^6×34^); (ii) the Y-Y distance (distance between Y219^5×58^ and Y326^7×53^); (iii) the RMSD of the conserved NPxxY motif (from N322^7×49^ to C327^7×54^) relative to the inactive structure 2RH1;^5^ and (iv) the RMSD of the PIF motif (I121^3×40^ and F282^6×44^) relative to the inactive structure 2RH1. ^5^

#### Ionic Lock Distance

The ionic lock distance is defined as the distance between the C*_α_* atoms of R131^3×50^ and L272^6×34^. The receptor is considered to be in an inactive state if this microswitch value is below 1.05 nm; it is considered to be active if the value is greater than 1.2 nm. This distance correlates to a salt bridge between R131^3×50^ and E268^6×30^, which is often observed in the inactive state of Class A GPCRs.

#### Y-Y Distance

The Y-Y distance is defined as the distance between the C*_ζ_* atoms of Y219^5×58^ and Y326^7×53^. The receptor is considered in an inactive state if this microswitch value is above 1.46 nm, while it is considered active if it is smaller than 0.8 nm. Due to the position of the residues that define this microswitch within the receptor, it is typically associated with activation. The involved tyrosines are considered to play a gating role, favoring or impeding ion access to the ion binding site.

#### NPxxY RMSD

The NPxxY RMSD microswitch is defined as fitting the structure of the receptor to the crystallographic inactive state structure (2RH1), considering one subset of the C*_α_* atoms of the transmembrane portion, and computing the root mean square deviation of the heavy atoms in the backbone of the residues from N322^7×49^ to C327^7×54^. The receptor is considered to be in the inactive state if the RMSD is less than 0.2 nm and active if the RMSD is greater than 0.34 nm. This microswitch detects the rotation of the intracellular portion of TM7.

#### PIF RMSD

The PIF RMSD microswitch is defined as fitting the structure of the receptor to the crystallographic inactive state structure (2RH1), considering one subset of the C*_α_* atoms of the transmembrane portion of the receptor, and computing the root mean square deviation of the heavy atoms of the residues I121^3×40^ and F282^6×44^. The receptor is considered to be in the inactive state if the RMSD is less than 0.22 nm and active if this RMSD is greater than 0.31 nm.

## Results and discussion

### Spontaneous inactivation of *β*_2_AR

Our initial aim was to observe the transition from the experimental determined active-state conformation to the inactive one through unbiased MD simulations at different pH values. Considering the available literature regarding *in silico* calculations of such conformational change by means of classical force fields,^10–12^ we estimated that a simulation length of 1 *µ*s would have been sufficient to observe this phenomenon. To achieve reliable statistical significance, we ran five independent replicates for each of the six pH values studied (from 4 to 9).

We observed a spontaneous inactivation of the *β*_2_AR receptor with a pH-dependent behavior, as expected from experimental evidence.^13^ First, we computed the C*_α_* RMSD of the transmembrane residues of *β*_2_AR, using the experimental structures of the active (3P0G) and inactive (2RH1) states as a reference. The RMSD from the inactive structure becomes almost immediately smaller than the RMSD from the active structures for the majority of the replicates at pH values above 5 (see Figure S3).

To investigate the transition in more detail, we evaluated the molecular microswitches that are usually considered a proxy for the activation state of *β*_2_AR, as defined in the Methods section (the ionic lock formation, the Y-Y distance, the PIF and the NPxxY motifs formation). A graphical description of the microswitches and the average behavior observed in the trajectories per pH value are in Figure 1 (the timeseries for all the microswitches are available in the Figures S5-8, while a recap scheme is in Figure S4 in the Supporting Information).

**Figure 1:**
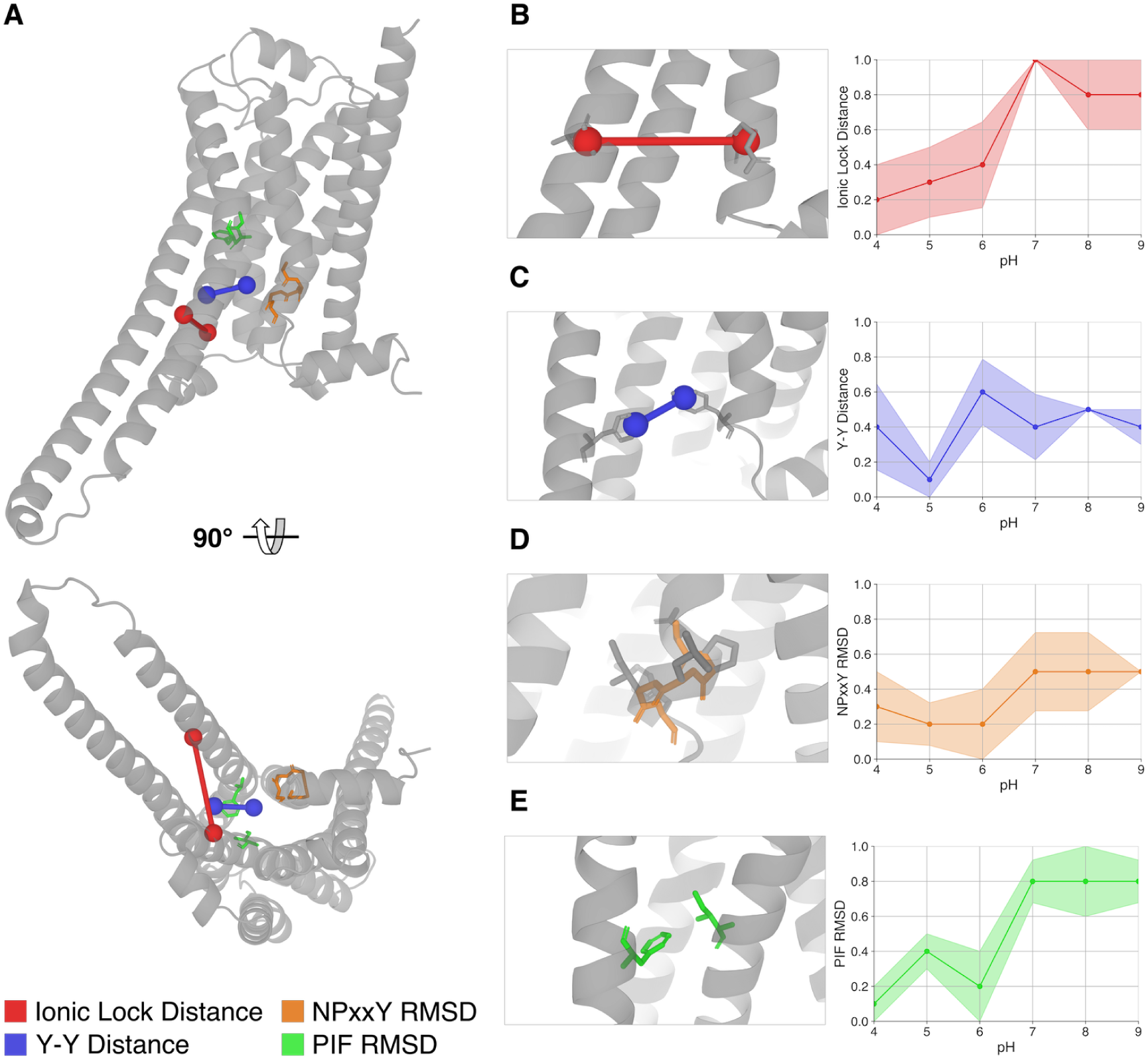
Microswitches considered as a proxy of the *β*_2_AR activation state in simulations. A) Position of the microswitches in the structure of the *β*_2_AR receptor. On the right side of the figure a zoom-in on the residues (represented in sticks) that define the four microswitches and their behavior as a function of the pH value: B) is the ionic lock distance (in red), C) the Y-Y distance (in blue), D) the NPxxY RMSD (in orange), and E) the PIF RMSD (in green). Each plot represents the average microswitch activation state observed over the five replicates per pH condition: the value 1 corresponds to the microswitch indicating the inactivation state (according to the threshold defined in the methods section), the value 0 corresponds to activation, while 0.5 indicates a state between the two thresholds. The error shown is the standard deviation of the mean.

As Figure 1 shows, there is no complete agreement among the microswitches. The ionic lock distance, PIF and NPxxY RMSD exhibit sigmoid properties (although NPxxY never reaches the fully inactivated threshold). Conversely, the Y-Y distance does not appear to follow a pH-dependent behavior: we observe a different orientation of the two residues that does not align with the threshold specified in the microswitch definition.^6,12,45^ This apparent discrepancy between Y-Y and the other microswitches may be attributed to the fact that such distance depends on slow water movement, which occurs on longer timescales ^46^ (up to the ms range). Despite this inconsistency in the activation state of some of the microswitches, visual inspection reveals the inward tilting and stabilization of TM6 in all the simulations with a pH greater than 5. This rearrangement corresponds with the formation of the ionic lock microswitch and leads to the closure of the binding interface of the G protein. There is also a general stabilization of a conformation which is closer to the inactive state.

### Dynamics of protonation states in titratable sites

Following the microswitch analysis, we took advantage of the constant-pH model to study the variable protonation state of titrable residues. From a mechanistic point of view, the pH change and the consequent charge movement (both for the protonation state and ionic movement) can trigger conformational rearrangements in protein structures. Focusing on *β*_2_AR, the most studied residue in this regard is D79^2×50^, where its protonation state has been considered a fundamental key element in the activation process of this class A GPCR.^10,11,19,32^ In our simulations we analyzed the population of protonated/deprotonated titrable residues in function of the pH. We separately analyzed acidic residues, basic residues and histidines considering the *λ* value as the proxy of protonation state. Following the prescription of the method^29,30^ a residue with *λ <* 0.2 is considered neutral while with *λ >* 0.8 is considered charged.

Starting from the acidic group of residues in *β*_2_AR, we show the results in Figure 2. Given the pK*_a_* values of the carboxylic acid side chains of aspartic acid (3.9) and glutamic acid (4.1), we expected that solvent-exposed acidic residues would exhibit intermediate protonation states at pH 4, and be predominant deprotonated (negatively charged) at higher pH values. This hypothesis was confirmed for all the aspartic and glutamic acid residues located in the intracellular loops (ICLs), extracellular loops (ECLs), and helix 8 (H8). This observation suggests potential electrostatic interactions with sodium ions, which are discussed in the following section.

**Figure 2:**
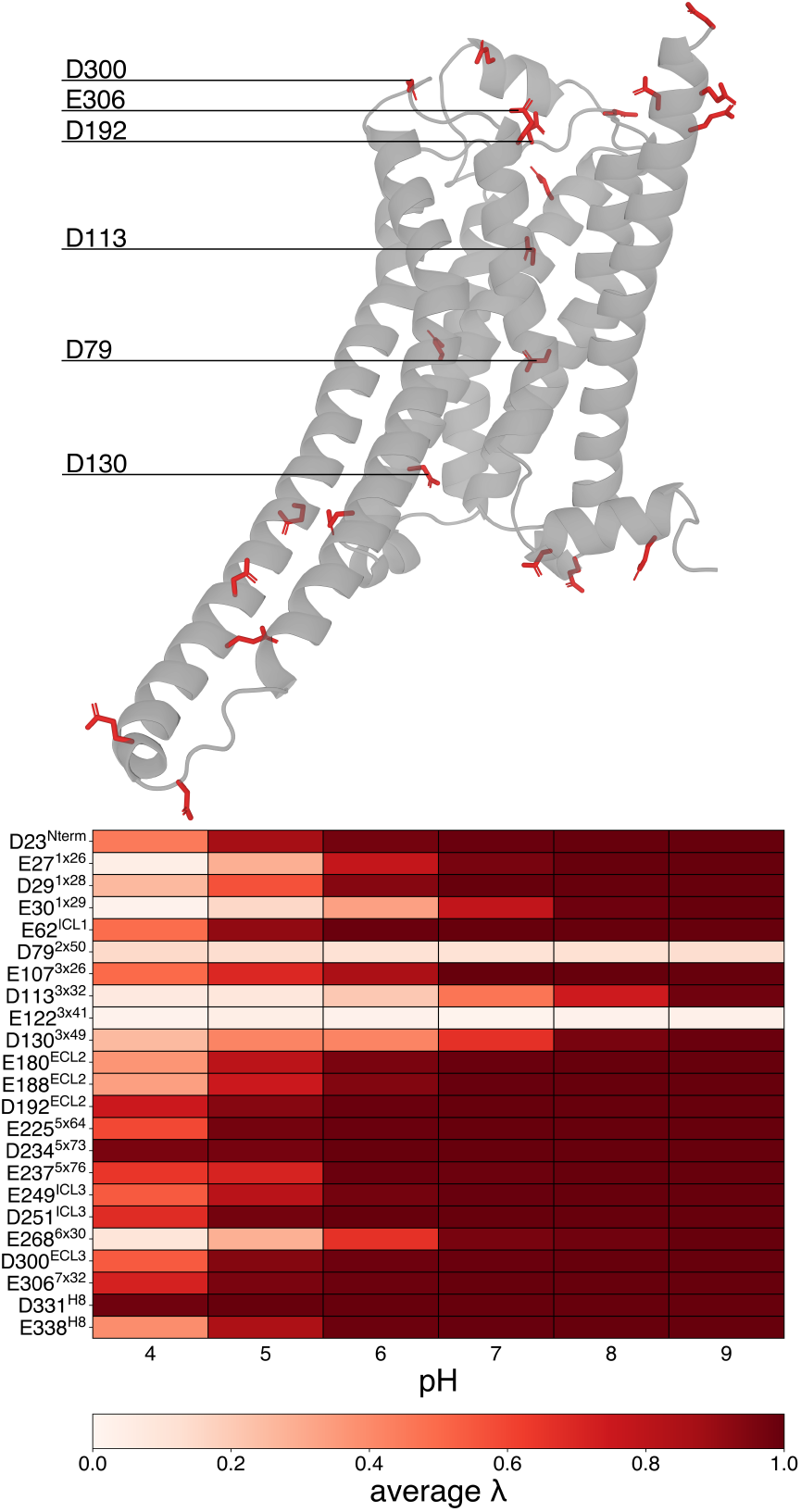
Titrable acidic residues and their protonation state at different pH. On top, the structure of *β*_2_AR with the glutamic and aspartic acid residues highlighted in red sticks. Below, a colormap representing the protonation state of all the acidic residues for different pH values (*λ* = 1 means charged −1, *λ* = 0 means neutral), averaged over five replicas.

We focused on residues whose behavior deviates from that of fully solvated acidic residues, and those that appear to undergo a protonation change between pH 5 and 6. This pH range corresponds to the lower limit at which *β*_2_AR inactivation can still be observed. The atypical protonation behavior of these residues led us to hypothesize their potential involvement in the inactivation transition.

E27^1×26^ and E30^1×29^ are close together, mutually causing a shift in the pK*_a_* value. Considering the third residue in that vicinity, D29^1×28^, there is a complete transition to the charged state at a pH between 5 and 6, which can be connected to interactions with sodium ions on the extracellular side of the receptor. E122^3×41^ remains protonated (neutral) across all pH conditions in the constant-pH simulations, likely due to its exposure to the hydrophobic portion of the membrane. D130^3×49^ is located on the intracellular side of the receptor, at a possible entry point for sodium ions into the ion binding pocket. Its partially buried position likely justifies the small observed pK*_a_* shift. E268^6×30^ is the acidic residue which forms the salt bridge with R131^3×50^ in the ionic lock microswitch. As described in the previous section, the ionic lock forms under conditions favoring the inactive state. This supports a deprotonated state of E268^6×30^, consistent with its role in stabilizing the salt bridge. D113^3×32^ is known to be a key residue for ligand binding via electrostatic interactions. ^47^ In our simulations, despite being in the middle of the membrane, it remains fully hydrated. Starting at pH 7, we observed multiple protonation/deprotonation transitions on such residue, which are likely driven by the interaction with sodium ions, as previously reported in classical MD simulations.^21^ The last and most interesting residue that deviates from the the expected pK*_a_*behavior is D79^2×50^, which is never observed in its deprotonated (charged) state, regardless of the pH. This contrasts with previous computational studies, which consider the negatively charged state a necessary condition to trigger the inactivation of *β*_2_AR,^11,19,20^ here observed from pH 6 onward. The absence of the charged form of D79^2×50^ strongly suggests a lack of interaction with positively charged groups, hinting towards the absence of sodium ions in the pocket.

Moving to the analysis of the basic residues, we observe that none of them undergo a clear protonation switch from the neutral to the charged state across the range of pH values studied in this work. Only a few residues showed slight changes in their *λ* values at pH 9. Due to the lack of substantial pH-dependent behavior, the full analysis is provided in the Supporting Information file (Figure S9).

Histidines are treated as multi-protonation state residues in the constant-pH implementation we used. In practice, these residues are modeled as a three-state system, including two neutral forms (*δ*- and *ε*-protonated) and the double protonated charged form. In our analysis, we focused only on the charged versus neutral distinction. The protonation behavior of histidine residues is summarized in Figure 3.

**Figure 3:**
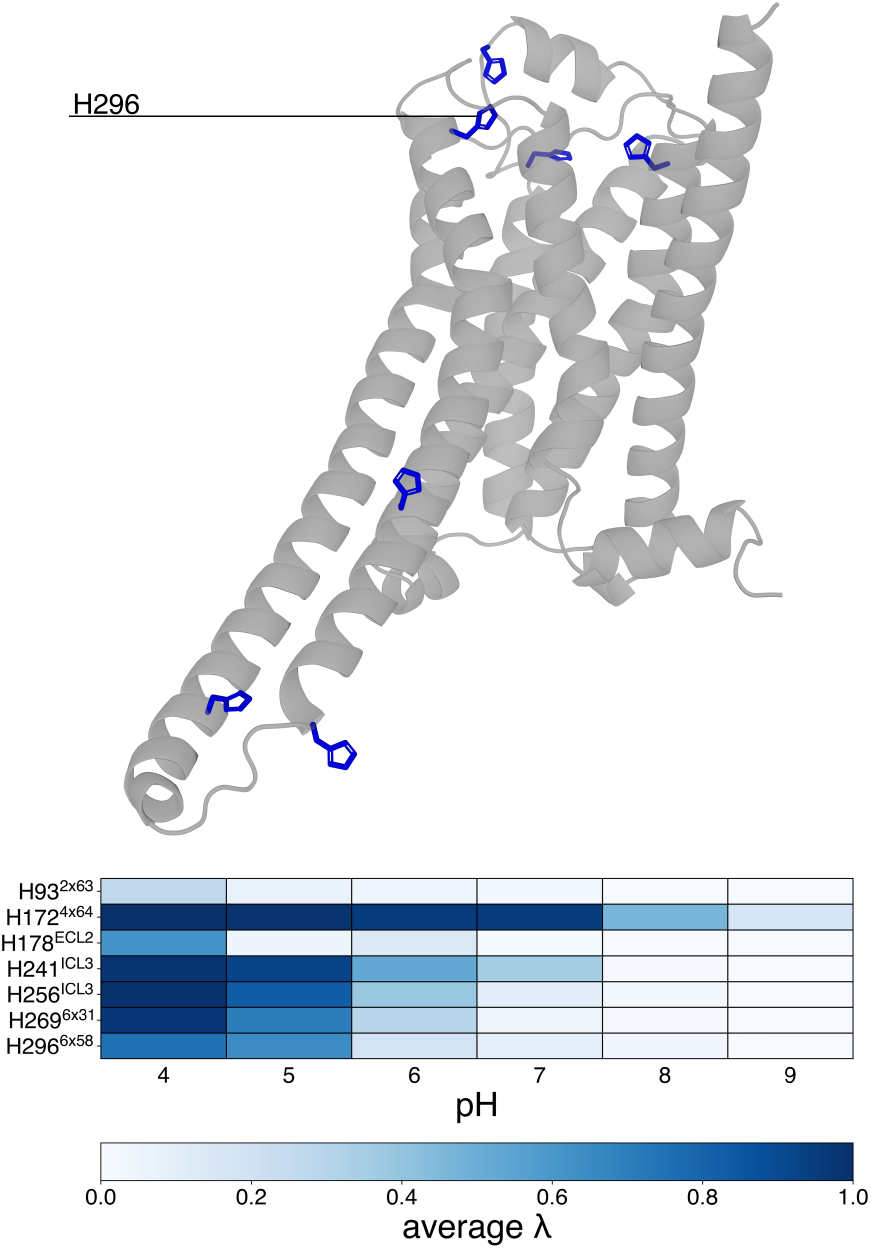
Titrable histidine residues and their protonation state at different pH. On top, the structure of *β*_2_AR with histidine residues highlighted in blue sticks. Below, a colormap representing the charge state of the histidine residues (*λ* = 1 means charged +1, *λ* = 0 means neutral). The *λ* value is averaged over five replicates per pH condition.

Interestingly, the protonation switch from the charged to the neutral state among histidine residues can be roughly divided into three groups, based on the pH range in which the transition occurs: (i) between pH 5 and 6 for H241^ICL3^, H256^ICL3^, H269^6×31^, and H296^6×58^, (ii) between pH 7 and pH 8 for H172^4×64^, and (iii) those that do not show a predominant charged state at any pH value. Considering the first group, we do not expect a structural role in receptor inactivation for the ICL3 residues, as this intracellular loop undergoes significant conformational change in the absence of a binding partner. H269^6×31^ is located on the intracellular side of the receptor and mainly interacts with the membrane. The most interesting residue is H296^6×58^, which is positioned close to D300^ECL3^, at the entrance of the intramembrane portion of the receptor accessible for solvent and ions. This location suggests a possible role for this residue in ion trafficking.

The protonation behavior of H172^4×64^, the sole member of the second group, can be explained by its proximity to E107^3×26^, which stabilizes its charged state up to pH 7. Lastly, H93^2×63^ and H178^ECL2^ never display a charged state for most of the simulation time at any pH level. This behavior is likely due to their constant exposure to the solvent on the extracellular side of the receptor.

### Dynamics of Na^+^ ions in the inactivation process

To investigate the behavior of sodium ions in proximity to residues D79^2×50^ and D113^3×32^, we performed a systematic analysis of the MD trajectories. For each replica and pH condition, we computed the distance between the C*_γ_* atom of either D79^2×50^ or D113^3×32^ and each sodium ion in solution. For D113^3×32^, a binding event was defined as any frame in which a sodium ion approached within 5 Å of the residue. In contrast, due to the lack of closer approaches, a more permissive distance threshold of 10 Å was applied to D79^2×50^. Ions that satisfied these distance criteria were identified and analyzed further in terms of their time-dependent distance profiles and permanence, which is defined as the number of frames spent within the specified cutoff. Additionally, for the five replicas at pH 7, we examined the trajectories of these ions to characterize their pathways to highlight the differences between constant-pH simulations and the fixed protonation ones available in the literature. ^21^ A graphical representation of the portion of the receptor visited during pH 7 simulations by sodium ions is in Figure 4.

**Figure 4:**
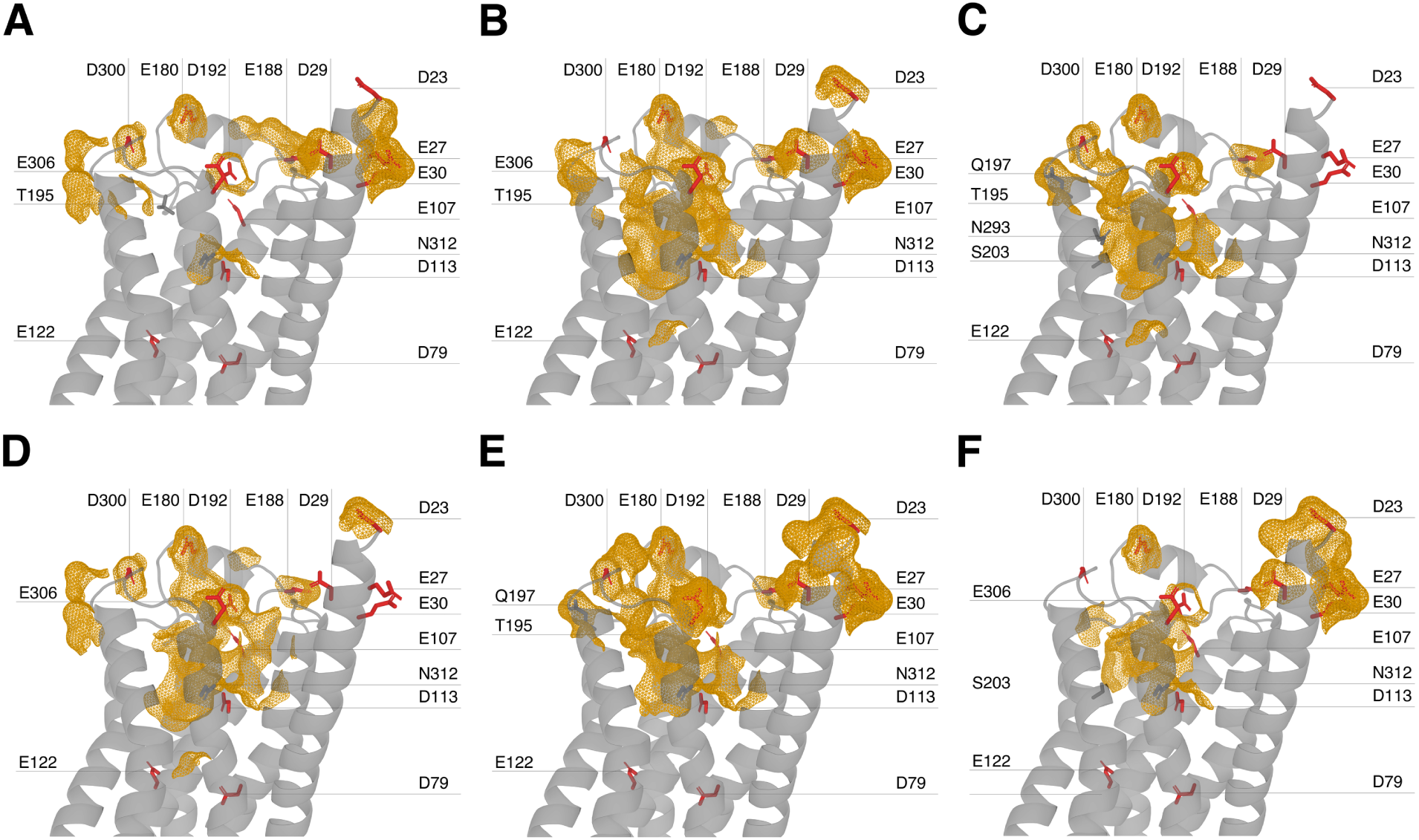
Sodium ion binding pathways in pH 7 simulations across five MD replicas. The panels (A–F) show the portions of the receptor visited by Na^+^ ions during binding events in five independent molecular dynamics simulations at pH 7. Red sticks represent Asp and Glu side chains (mainchain atoms and hydrogens not shown), visualized independently of Na^+^ interaction. Gray sticks indicate polar side chain residues within 5 Å of Na^+^ during its approach to residue D113^3×32^. Orange mesh surfaces highlight regions where Na^+^ was within 3.5 Å of surrounding atoms for at least 5 frames along the trajectory. Panels correspond to: A) replica 1, binding event 1; B) replica 1, binding event 2; C) replica 2; D) replica 3; E) replica 4; and F) replica 5.

Considering the available literature on the role of the ion binding pocket in *β*_2_AR, the main observation of this analysis is that, regardless of the pH value, there are no sodium ions interacting with residue D79^2×50^ at any point in the inactivation process.

Although the specific pathways vary between replicas, the recurrence of certain residues suggests that they may define preferential routes or act as key gating points during sodium ion entry and stabilization. A full description of all sodium ion pathways is available in the SI. Across pH conditions, the closest approaches to D79^2×50^ ranged from 8.1 Å (replica 2, pH 7) to 9.9 Å (replica 5, pH 8); however, these consistently coincided with direct interactions involving D113^3×32^. This residue showed no ion contacts at pH 4–5 and only sporadic, short-lived events at pH 6. In contrast, at pH 7, all five replicas exhibited multiple binding events: two in replica 1, an exceptionally stable interaction in replica 2 (∼826.6 ns), and long-lived contacts in replica 3 (∼482 ns), replica 4 and replica 5. At higher pH values (8 and 9) both the frequency and duration of binding events increased, with up to ten independent contacts per replica and residence times ranging from sub-nanosecond transients to interactions lasting over 900 ns.

Focusing on the trajectories at pH 7, pathway analysis revealed that sodium ions navigate a conserved corridor of polar and acidic side chains—chiefly D300^ECL3^, D192^ECL2^, T195^ECL2^, Q197^5×37^, N293^6×55^, N312^7×38^, and S203^5×43^—before stabilizing at D113^3×32^. These residues consistently emerge as key gating points that guide and coordinate sodium entry into the allosteric site.

Furthermore, none of the simulations showed sodium ions overcoming the hydrophobic core consisting of M82^2×53^, V117^3×36^, F208^5×47^, W286^6×48^, and the PIF motif residues I121^3×40^ and F282^6×44^. This region may form an effective barrier that prevents ions from accessing D79^2×50^.

In comparison, Wang et al.^21^ found that, in simulations involving deprotonated D79^2×50^, Na^+^ ions progressed from an initial engagement at D113^3×32^, through coordination with N312^7×38^, into the conserved ion binding pocket close to D79^2×50^, where they were further coordinated by S319^7×46^ and F282^6×44^. Crucially, when D79^2×50^ was protonated in their simulations, sodium ions no longer penetrated into the binding pocket, indicating a lack of electrostatic driving force. This outcome closely mirrors our constant-pH simulations at neutral pH, in which ions strongly bind to D113^3×32^ but do not proceed into the ion binding site. Considering the intracellular part of the receptor, D130^3×49^ is the only acidic residue in direct contact with the solvent. We performed the same distance-based analysis for all sodium ions and observed only extremely brief binding events, limited to simulations at pH 6 and pH 7. The corresponding time series are shown in Figure S11 of the Supporting Information.

In summary, our results suggest that the dynamics of sodium ions influence receptor mobility during the inactivation process. As expected, low pH conditions (4-5) do not favor the binding of positively charged sodium ions to acidic residues. This effect is likely due to the protonation of H296^6×58^, which remains positively charged up to pH 5, hindering the passage of Na^+^ ions through the gate formed by D300^ECL3^ that leads to the transmembrane part of the receptor. At intermediate pH values (6-7), we observe more stable and durable interactions between sodium ions and residues in the transmembrane portion of the receptor. At higher pH levels (8-9), sodium mobility increases, resulting in more frequent but transient binding events with acidic side chains. These observations support the hypothesis that sodium ion binding is most stable around physiological pH (∼7), where it may facilitate the progression of the inactivation process. In contrast, crucial interactions, such as the ionic lock, do not form at lower pH. At higher pH, the receptor can still inactivate, but with a less coordinated sequence of binding events, potentially slowing down the global conformational transition. For completeness, we also analyzed the binding of chloride ions to positively charged residues during the simulations. No stable interactions were detected between Cl^-^ ions and the protein. A quantitative analysis is provided in Figure S12 of the Supporting Information.

## Conclusion

In this work, we present a systematic investigation of the inactivation of the *β*_2_AR using constant-pH MD simulations. Our results support the experimental observed pH-dependent nature of the inactivation process,^13^ which is closely linked to protonation changes of acidic side chains. One of the main residues is E268^6×30^, which is involved in the formation of the ionic lock, a fundamental microswitch of the inactivation process.

We note, however, that while our 1 *µ*s-long replicas are relatively extensive, they may still undersample slow, ms-scale hydration and gating events around the NPxxY motif. Future work with longer runs will be needed to fully capture this critical water-mediated transition. One of the most surprising findings of this study is the absence of Na^+^ ions in the ion binding pocket near D79^2×50^ and S120^3×39^ under all simulation conditions tested. Despite this, the receptor undergoes the conformational transitions associated with the inactive state. This contrasts with the majority of previous computational works^11,19–21^ that focused on the role of sodium ions and the protonation of D79^2×50^, concluding that a deprotonated state of the latter and the binding of sodium ions were key steps toward inactivation of the receptor. However, in our simulations we observe an unperturbed neutral state of D79^2×50^, and a stable ion binding position in D113^3×32^, hampering any deeper penetration of Na^+^ ions into the conserved ion-binding pocket. Notably, this behavior is consistent with observations reported in the work of Wang *et al.*,^21^ where the protonated D79^2×50^ similarly prevented sodium insertion into the ion-binding site.

Experimental evidence supporting the presence of Na^+^ near D79^2×50^ in *β*_2_AR is described by Katritch *et al.* in their work on the role of sodium in GPCR signaling:^15^ in a crystal structure of *β*_2_AR (PDB ID: 2RH1), electron density is observed near D79^2×50^ that is consistent with a bound Na^+^, although the resolution is insufficient to unambiguously assign the ion. However, the same structure lacks the ionic lock, a feature expected according to functional mutagenesis studies,^48,49^ which showed that *β*_2_AR is constitutively active if D130^3×50^ and/or R131^6×30^ are mutated. We propose that these discrepancies may arise due to the presence of strain in the crystal of *β*_2_AR, which could trap the receptor in a local conformational minimum which does not correspond to the global free energy minimum of the system, as already shown for other systems.^50,51^ Finally, we also examined the possibility of stable chloride ion binding at any pH, but found no such interactions. This supports the hypothesis that modulation of *β*_2_AR by anions occurs only via interactions mediated by the G protein.^52,53^

These simulations employ the fixed-charge CHARMM36m/TIP3P force field, which omits explicit electronic polarization, potentially crucial for modeling free energy barriers.^54,55^ Recent polarizable constant-pH MD implementations have demonstrated improved pK*_a_*predictions and deeper insights into electrostatic coupling, ^56^ and represent a promising direction for refining our mechanistic picture.

More broadly, this work demonstrates the power of constant-pH MD not only for *β*_2_AR but for any pH-sensitive GPCR, which are currently an extremely active research topic.^57,58^ Finally, the atomistic insights into protonation-driven inactivation and Na^+^ coordination extend to a broader understanding of ions role in other GPCRs,^59^ and it may guide the design of pH-selective agonists or allosteric modulators with improved therapeutic profiles. In conclusion, our findings underscore the need for the community to adopt increasingly accurate electrostatic models to faithfully represent such interactions. The constant-pH MD framework proves extremely useful in contexts where pH and electrostatics dominate the behavior of biomolecules.

## Supporting information

Supplemental tables, figures, and full description of Na+ binding pathways

## Data Availability Statement

All the input file to reproduce this results and the trajectories of the runs shown here are available on the Zenodo repository with the DOI 10.5281/zenodo.15744221.

## Acknowledgement

The authors thank Mercedes Alfonso-Prieto and Marina Casiraghi for useful discussion. R.C. was supported by Università degli Studi di Milano (Piano di supporto alla ricerca di ateneo Linea 2 2025-DBS Capelli). F.B. acknowledges the Next Generation EU project PRIN 2022 (20225X2FS5, CUP B53D23015140006).

## Supporting Information Available

Extended description of the Na^+^ ions pathways observed during MD simulations, tables with the Ballesteros-Weinstein numbering of *β*_2_AR, with the main events observed in every MD replica. Snake diagram of the *β*_2_AR with highlighted titrable residues, timeseries of RMSD, microswitches, Na^+^ ions binding, and heatmap of basic residues protonation states and Cl^-^ ions binding to histidines.

## TOC Graphic

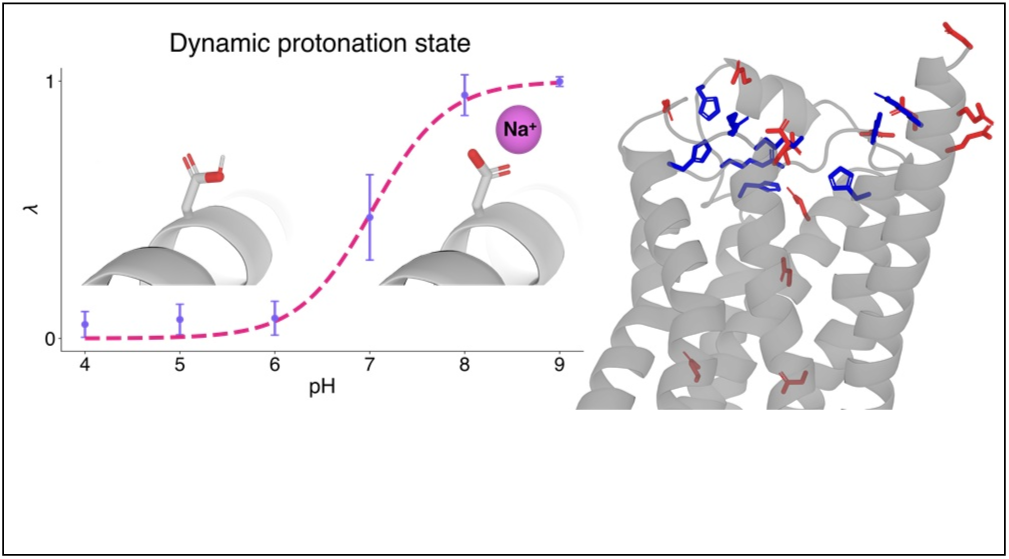

