## Supplemental tables, figures, and full description of Na+ binding pathways for "Constant-pH simulation of the human *β*_2_ adrenergic receptor inactivation"

### Supporting Information for: "Constant-pH simulation of the human $\beta_2$ adrenergic receptor inactivation"

#### Description of the $Na^+$ binding pathways

We performed a systematic analysis of the MD trajectories using the MDAnalysis library (version 2.7),[1] focusing on the $Na^+$ ions which moved close to residues $D79^{2x50}$ and $D113^{3x32}$. In replica 2 at pH 7, a single ion was observed to reach a minimum distance from $D79^{2x50}$ of 8.1 Å; in replica 3 at pH 8, the minimum distance was 9.4 Å; in replica 5 at pH 8, the minimum distance was 9.9 Å; in replica 5 at pH 9, the minimum distance was 8.9 Å. Analysis of the trajectories showed that these minimum distances corresponded to interactions with residue $D113^{3x32}$. Regarding the latter residue $D113^{3x32}$, no sodium ions approached within 5 Å at pH 4 or pH 5. Interactions were observed from pH 6 onward.

Considering the sodium entrance in every simulation, we have that for pH 6, a single binding event lasting 16.4 ns occurred at the end of replica 1 (distance plot S10A). In replica 4, three sodium ions interacting with the residue were observed at different times for a total of 6.5 ns (distance plot S10B). Interestingly, the ion in replica 1 also interacted with residues E180$^{ECL2}$ and D300$^{ECL3}$ in the interval between ∼622.5 ns and ∼655.5 ns (distance plot A). At pH 7, residue D113$^{3x32}$ is engaged by ions in all five replicas. In replica 1, the first binding event occurred at 333.6 ns and lasts 1.8 ns. A different sodium ion entered at 431.7 ns and remained for  84.9 ns (distance plot S10C). The most persistent event occurred in replica 2, where the ion remained within 5 Å of the residue for  826.6 ns (distance plot S10D). Similarly, replica 3 showed a highly stable interaction, with the ion maintaining contact for 482 ns (distance plot S10E), while replicas 4 and 5 displayed shorter contacts of 15.2 ns and 4.9 ns (distance plots S10F and S10G), respectively. At pH 8, differently from pH 7, the frequency of multiple independent binding events for a single replica increased: nine ion-residue interactions in replica 3 and ten in replica 5. In detail, replica 1 featured three ions contacting the residue from 82.6 to 484.0 ns (distance plot S10H). Replica 2 showed a single, highly stable event (the longest observed here), with the ion maintaining the interaction for 920.7 ns (distance plot S10I). Replica 3 displayed a broad diversity of transient and stable events, with interaction durations varying from  0.1 ns to 102.7 ns (distance plot S10J). Replica 4 highlighted another long event, with the ion bound to the residue for 431.3 ns (distance plot S10K). Replica 5 contributed additional interactions, ranging from 3.3 ns to 85.2 ns (distance plot S10L). At pH 9, interaction events with residue 3x32 persisted across all replicas. Notably, only replica 1 exhibited a single binding event, which is the second most stable in the dataset, lasting 859.4 ns (distance plot S10M). Replica 2 featured multiple ions interacting at staggered times (distance plot S10N). Replica 3 included a highly persistent contact of 739.4 ns (distance plot S10O), while replica 4 showed both transient and stable events, including a  101.8 ns contact (distance plot S10P). Finally, replica 5 was marked by a particularly long interaction with the ion that remains within 5 Å of residue D113$^{3x32}$ for over 519.1 ns (distance plot S10Q).

According to the microswitches analysis (see the previous section), the pH 7 systems showed a better agreement in the majority of the features considered indicative of receptor inactivation during the simulation. Based on this observation, we focused our attention on the trajectories at pH 7, paying particular attention to the paths taken by sodium ions towards residue D113$^{3x32}$. We analyzed the sequence of polar and negatively charged residues encountered along the trajectory of each ion that reached D113$^{3x32}$ within 5 Å. These residues were defined as those interacting within 5 Å of the ion. Two independent ion binding events were identified in replica 1. During the first event, the ion first engaged with residue D300$^{ECL3}$, then moved through T195$^{ECL2}$, before being coordinated by both D113$^{3x32}$ and N312$^{7x38}$. As previously described, this interaction was relatively short-lived, lasting around 1.8 ns. The second event in the same replica was notably longer. In this case, a different ion interacted with D192$^{ECL2}$ before continuing through the extracellular cavity via T195$^{ECL2}$, and ultimately reached D113$^{3x32}$ and N312$^{7x38}$. In replica 2, a single ion enters via D300$^{ECL3}$ and Q197$^{5x37}$. It then proceeded through T195$^{ECL2}$ to N293$^{6x55}$ and S203$^{5x43}$ before finally reaching D113$^{3x32}$. In Replica 3, the ion path involved initial contacts with E306$^{7x32}$ and D192$^{ECL2}$, followed by coordination with N312$^{7x38}$ before engaging with D113$^{3x32}$. Replica 4 presented a more complex trajectory. First, the ion interacted with D192$^{ECL2}$, then with Q197$^{5x37}$. Subsequently, it engaged with T195$^{ECL2}$ before returning to D192$^{ECL2}$ and D300$^{ECL3}$. Finally, it stabilized at D113$^{3x32}$ assisted by N312$^{7x38}$. In replica 5, the ion entered through D300$^{ECL3}$, D192$^{ECL2}$, and moved to D113$^{3x32}$ and N312$^{7x38}$, eventually contacting S203$^{5x43}$. These observations suggested that the approach of ions to D113$^{3x32}$ involves consistent coordination by acidic and polar residues, such as D300$^{ECL3}$, D192$^{ECL2}$, and T195$^{ECL2}$.

Table S1: Ballesteros-Weinstein generic numbering for human $\beta_2$-adrenoceptor, as listed in the GPCRdb[2] (https://gpcrdb.org/residue/residuetabledisplay, accessed on December 2024).

| **TM1** | | **TM2** | | **TM3** | | **TM4** | | **TM5** | | **TM6** | | **TM7** | |
|---|---|---|---|---|---|---|---|---|---|---|---|---|---|
| 1x25 | Q26 | 2x37 | T66 | 3x21 | G102 | 4x38 | T146 | 5x36 | N196 | 6x24 | S262 | 7x30 | R304 |
| 1x26 | E27 | 2x38 | V67 | 3x22 | N103 | 4x39 | K147 | 5x37 | Q197 | 6x25 | K263 | 7x31 | K305 |
| 1x27 | R28 | 2x39 | T68 | 3x23 | F104 | 4x40 | N148 | 5x38 | A198 | 6x26 | F264 | 7x32 | E306 |
| 1x28 | D29 | 2x40 | N69 | 3x24 | W105 | 4x41 | K149 | 5x39 | Y199 | 6x27 | C265 | 7x33 | V307 |
| 1x29 | E30 | 2x41 | Y70 | 3x25 | C106 | 4x42 | A150 | 5x40 | A200 | 6x28 | L266 | 7x34 | Y308 |
| 1x30 | V31 | 2x42 | F71 | 3x26 | E107 | 4x43 | R151 | 5x41 | I201 | 6x29 | K267 | 7x35 | I309 |
| 1x31 | W32 | 2x43 | I72 | 3x27 | F108 | 4x44 | V152 | 5x42 | A202 | 6x30 | E268 | 7x36 | L310 |
| 1x32 | V33 | 2x44 | T73 | 3x28 | W109 | 4x45 | I153 | 5x43 | S203 | 6x31 | H269 | 7x37 | L311 |
| 1x33 | V34 | 2x45 | S74 | 3x29 | T110 | 4x46 | I154 | 5x44 | S204 | 6x32 | K270 | 7x38 | N312 |
| 1x34 | G35 | 2x46 | L75 | 3x30 | S111 | 4x47 | L155 | 5x45 | I205 | 6x33 | A271 | 7x39 | W313 |
| 1x35 | M36 | 2x47 | A76 | 3x31 | I112 | 4x48 | M156 | 5x46 | V206 | 6x34 | L272 | 7x40 | I314 |
| 1x36 | G37 | 2x48 | C77 | 3x32 | D113 | 4x49 | V157 | 5x461 | S207 | 6x35 | K273 | 7x41 | G315 |
| 1x37 | I38 | 2x49 | A78 | 3x33 | V114 | 4x50 | W158 | 5x47 | F208 | 6x36 | T274 | 7x42 | Y316 |
| 1x38 | V39 | 2x50 | D79 | 3x34 | L115 | 4x51 | I159 | 5x48 | Y209 | 6x37 | L275 | 7x43 | V317 |
| 1x39 | M40 | 2x51 | L80 | 3x35 | C116 | 4x52 | V160 | 5x49 | V210 | 6x38 | G276 | 7x45 | N318 |
| 1x40 | S41 | 2x52 | V81 | 3x36 | V117 | 4x53 | S161 | 5x50 | P211 | 6x39 | I277 | 7x46 | S319 |
| 1x41 | L42 | 2x53 | M82 | 3x37 | T118 | 4x54 | G162 | 5x51 | L212 | 6x40 | I278 | 7x47 | G320 |
| 1x42 | I43 | 2x54 | G83 | 3x38 | A119 | 4x55 | L163 | 5x52 | V213 | 6x41 | M279 | 7x48 | F321 |
| 1x43 | V44 | 2x55 | L84 | 3x39 | S120 | 4x56 | T164 | 5x53 | I214 | 6x42 | G280 | 7x49 | N322 |
| 1x44 | L45 | 2x551 | A85 | 3x40 | I121 | 4x57 | S165 | 5x54 | M215 | 6x43 | T281 | 7x50 | P323 |
| 1x45 | A46 | 2x56 | V86 | 3x41 | E122 | 4x58 | F166 | 5x55 | V216 | 6x44 | F282 | 7x51 | L324 |
| 1x46 | I47 | 2x57 | V87 | 3x42 | T123 | 4x59 | L167 | 5x56 | F217 | 6x45 | T283 | 7x52 | I325 |
| 1x47 | V48 | 2x58 | P88 | 3x43 | L124 | 4x60 | P168 | 5x57 | V218 | 6x46 | L284 | 7x53 | Y326 |
| 1x48 | F49 | 2x59 | F89 | 3x44 | C125 | 4x61 | I169 | 5x58 | Y219 | 6x47 | C285 | 7x54 | C327 |
| 1x49 | G50 | 2x60 | G90 | 3x45 | V126 | 4x62 | Q170 | 5x59 | S220 | 6x48 | W286 | 7x55 | R328 |
| 1x50 | N51 | 2x61 | A91 | 3x46 | I127 | 4x63 | M171 | 5x60 | R221 | 6x49 | L287 | | |
| 1x51 | V52 | 2x62 | A92 | 3x47 | A128 | 4x64 | H172 | 5x61 | V222 | 6x50 | P288 | | |
| 1x52 | L53 | 2x63 | H93 | 3x48 | V129 | | | 5x62 | F223 | 6x51 | F289 | | |
| 1x53 | V54 | 2x64 | I94 | 3x49 | D130 | | | 5x63 | Q224 | 6x52 | F290 | | |
| 1x54 | I55 | 2x65 | L95 | 3x50 | R131 | | | 5x64 | E225 | 6x53 | I291 | | |
| 1x55 | T56 | 2x66 | M96 | 3x51 | Y132 | | | 5x65 | A226 | 6x54 | V292 | | |
| 1x56 | A57 | 2x67 | K97 | 3x52 | F133 | | | 5x66 | K227 | 6x55 | N293 | | |
| 1x57 | I58 | | | 3x53 | A134 | | | 5x67 | R228 | 6x56 | I294 | | |
| 1x58 | A59 | | | 3x54 | I135 | | | 5x68 | Q229 | 6x57 | V295 | | |
| 1x59 | K60 | | | 3x55 | T136 | | | 5x69 | L230 | 6x58 | H296 | | |
| 1x60 | F61 | | | 3x56 | S137 | | | 5x70 | Q231 | 6x59 | V297 | | |
| | | | | | | | | 5x71 | K232 | 6x60 | I298 | | |
| | | | | | | | | 5x72 | I233 | 6x61 | Q299 | | |
| | | | | | | | | 5x73 | D234 | | | | |
| | | | | | | | | 5x74 | K235 | | | | |
| | | | | | | | | 5x75 | S236 | | | | |
| | | | | | | | | 5x76 | E237 | | | | |
| **ICL1** | | **ECL1** | | **ICL2** | | **ECL2** | | | | | | **H8** | |
| 12x48 | E62 | 23x49 | M98 | 34x50 | P138 | 45x50 | C191 | | | | | 8x47 | S329 |
| 12x49 | R63 | 23x50 | W99 | 34x51 | F139 | 45x51 | D192 | | | | | 8x48 | P330 |
| 12x50 | L64 | 23x51 | T100 | 34x52 | K140 | 45x52 | F193 | | | | | 8x49 | D331 |
| 12x51 | Q65 | 23x52 | F101 | 34x53 | Y141 | | | | | | | 8x50 | F332 |
| | | | | 34x54 | Q142 | | | | | | | 8x51 | R333 |
| | | | | 34x55 | S143 | | | | | | | 8x52 | I334 |
| | | | | 34x56 | L144 | | | | | | | 8x53 | A335 |
| | | | | 34x57 | L145 | | | | | | | 8x54 | F336 |
| | | | | | | | | | | | | 8x55 | Q337 |
| | | | | | | | | | | | | 8x56 | E338 |
| | | | | | | | | | | | | 8x57 | L339 |
| | | | | | | | | | | | | 8x58 | L340 |
| | | | | | | | | | | | | 8x59 | C341 |

Table S2: Sodium ions entries within 5 Å cutoff from D113$^{3x32}$. Each row corresponds to a single $Na^+$ ion observed in a specific replica and pH condition, with the ion identified by its atom ID and residue ID as in the reference structure protein_na.pdb (available in the Zotero repository). "Entry time" refers to the first frame in which the ion is observed within the cutoff, and "Length" indicates the total duration (in ns) the ion remained within 5 Å.

| pH | Replica ID | Ion Atom | Ion Residue | Entry time [ns] | Length [ns] |
|---|---|---|---|---|---|
| 6 | 1 | 5297 | 184 | 983.7 | 16.4 |
| 6 | 4 | 5303 | 190 | 366.9 | 0.7 |
| 6 | 4 | 5323 | 210 | 51.5 | 1.1 |
| 6 | 4 | 5330 | 217 | 13.1 | 4.7 |
| 7 | 1 | 5296 | 183 | 333.6 | 1.8 |
| 7 | 1 | 5323 | 210 | 431.7 | 84.9 |
| 7 | 2 | 5330 | 217 | 97.6 | 826.6 |
| 7 | 3 | 5305 | 192 | 67.4 | 482 |
| 7 | 4 | 5294 | 181 | 221.8 | 15.2 |
| 7 | 5 | 5290 | 177 | 241.7 | 4.9 |
| 8 | 1 | 5279 | 166 | 483.4 | 484 |
| 8 | 1 | 5291 | 178 | 79.3 | 118.1 |
| 8 | 1 | 5331 | 218 | 307.5 | 82.6 |
| 8 | 2 | 5295 | 182 | 61.8 | 920.7 |
| 8 | 3 | 5281 | 168 | 576.7 | 6.1 |
| 8 | 3 | 5300 | 187 | 140.1 | 102.7 |
| 8 | 3 | 5303 | 190 | 568.7 | 0.8 |
| 8 | 3 | 5304 | 191 | 675.5 | 2.6 |
| 8 | 3 | 5307 | 194 | 468.4 | 0.1 |
| 8 | 3 | 5308 | 195 | 251.2 | 2 |
| 8 | 3 | 5315 | 202 | 891.6 | 0.1 |
| 8 | 3 | 5317 | 204 | 698.8 | 21.7 |
| 8 | 3 | 5332 | 219 | 554.3 | 2.7 |
| 8 | 4 | 5317 | 204 | 535.1 | 431.3 |
| 8 | 5 | 5279 | 166 | 158.5 | 7.1 |
| 8 | 5 | 5282 | 169 | 743.6 | 1.3 |
| 8 | 5 | 5298 | 185 | 653.9 | 8.6 |
| 8 | 5 | 5299 | 186 | 417.1 | 4.4 |
| 8 | 5 | 5306 | 193 | 14.6 | 85.2 |
| 8 | 5 | 5310 | 197 | 189.2 | 22 |
| 8 | 5 | 5316 | 203 | 517.1 | 10.3 |
| 8 | 5 | 5320 | 207 | 566.3 | 5.8 |
| 8 | 5 | 5322 | 209 | 449.5 | 3.6 |
| 8 | 5 | 5325 | 212 | 489.6 | 3.3 |
| 9 | 1 | 5280 | 167 | 47.9 | 859.4 |
| 9 | 2 | 5284 | 171 | 267.0 | 10.5 |
| 9 | 2 | 5299 | 186 | 891.5 | 83.3 |
| 9 | 2 | 5302 | 189 | 418.2 | 106.3 |
| 9 | 2 | 5303 | 190 | 77.6 | 58.5 |
| 9 | 2 | 5308 | 195 | 622.6 | 169.6 |
| 9 | 2 | 5309 | 196 | 366.8 | 1.3 |
| 9 | 2 | 5314 | 201 | 156.2 | 82.8 |
| 9 | 2 | 5317 | 204 | 310.5 | 23.6 |
| 9 | 3 | 5281 | 168 | 917.0 | 7.8 |
| 9 | 3 | 5306 | 193 | 6.0 | 739.4 |
| 9 | 3 | 5312 | 199 | 938.2 | 47.8 |
| 9 | 4 | 5279 | 166 | 391.7 | 33.4 |
| 9 | 4 | 5282 | 169 | 290.6 | 6 |
| 9 | 4 | 5294 | 181 | 677.3 | 30.5 |
| 9 | 4 | 5307 | 194 | 925.8 | 4.4 |
| 9 | 4 | 5321 | 208 | 333.8 | 35.3 |
| 9 | 4 | 5324 | 211 | 750.6 | 97.5 |
| 9 | 4 | 5327 | 214 | 441.1 | 101.8 |
| 9 | 5 | 5316 | 203 | 112.2 | 519.1 |
| 9 | 5 | 5332 | 219 | 91.7 | 0.5 |

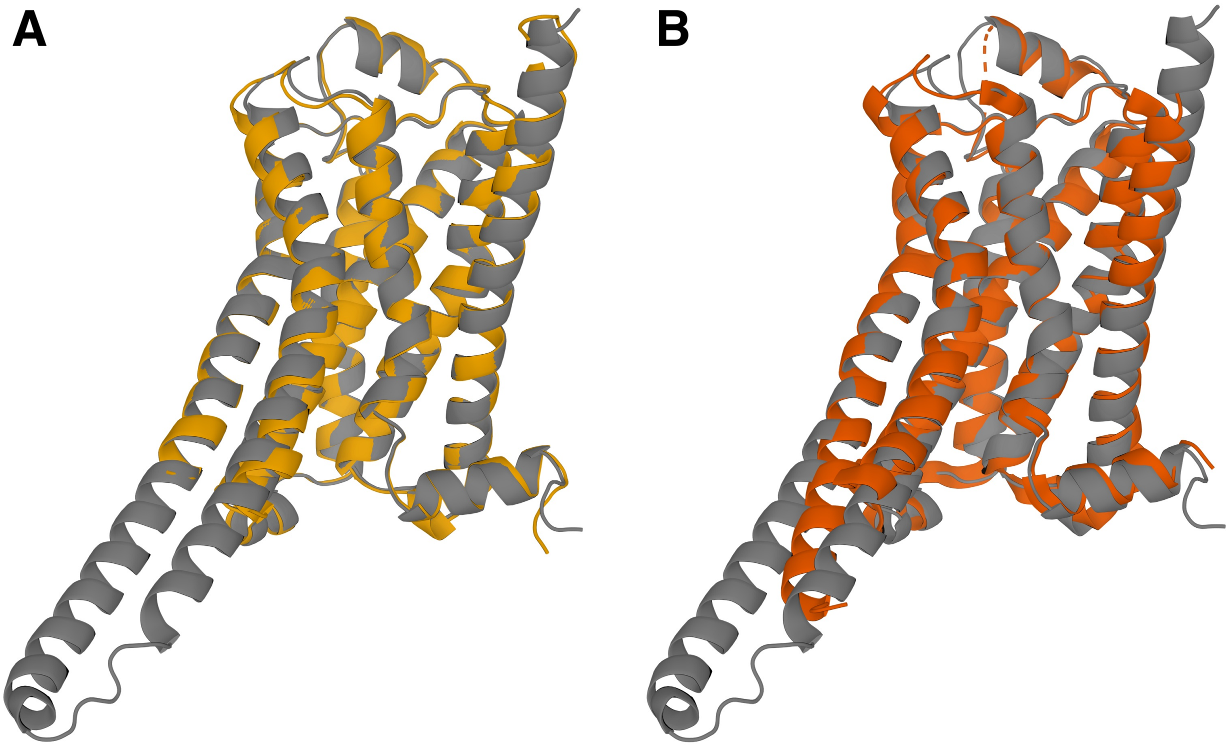


Figure S1: Comparison of the AlphaFold-reconstructed model and two experimental structures. A) Superposition between the model (gray) and the active (nanobody-bound) structure 3P0G (yellow). B) Superposition between the model (gray) and the active (G protein-bound) structure 3SN6 (orange). The initial model was obtained having as input the sequence of $\beta_2$AR and the Nb80 nanobody (residues 2-122), which explains the closer similarity to the nanobody-bound one.

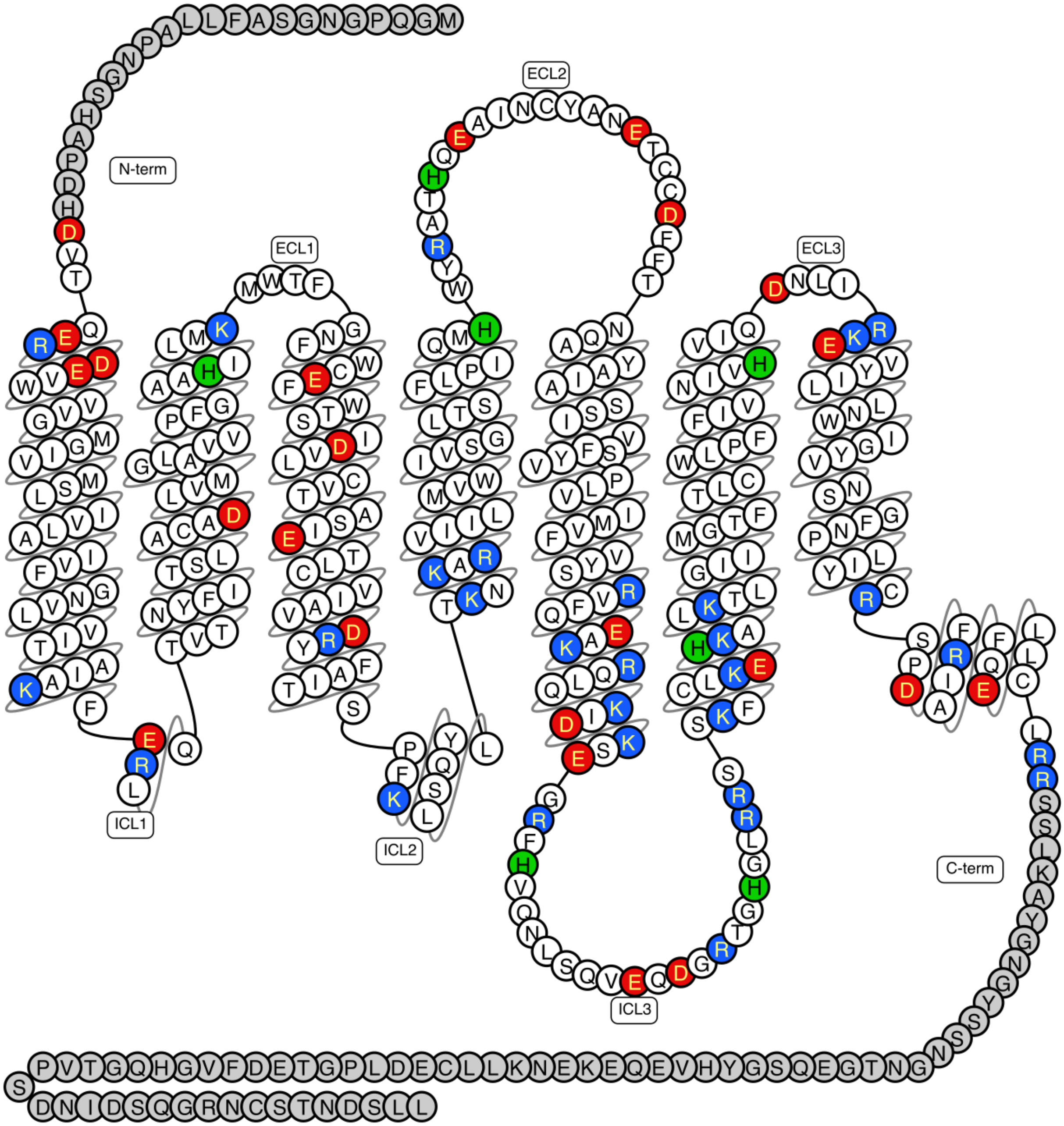


Figure S2: Snake diagram of the titrable residues in human $\beta_2$-adrenoceptor. The residues colored in red are the acidic residues, the blue ones are the basic ones, and the green one are histidines. The gray residues are the ones not included in our model. The snake diagram was obtained from the GPCRdb.[2]

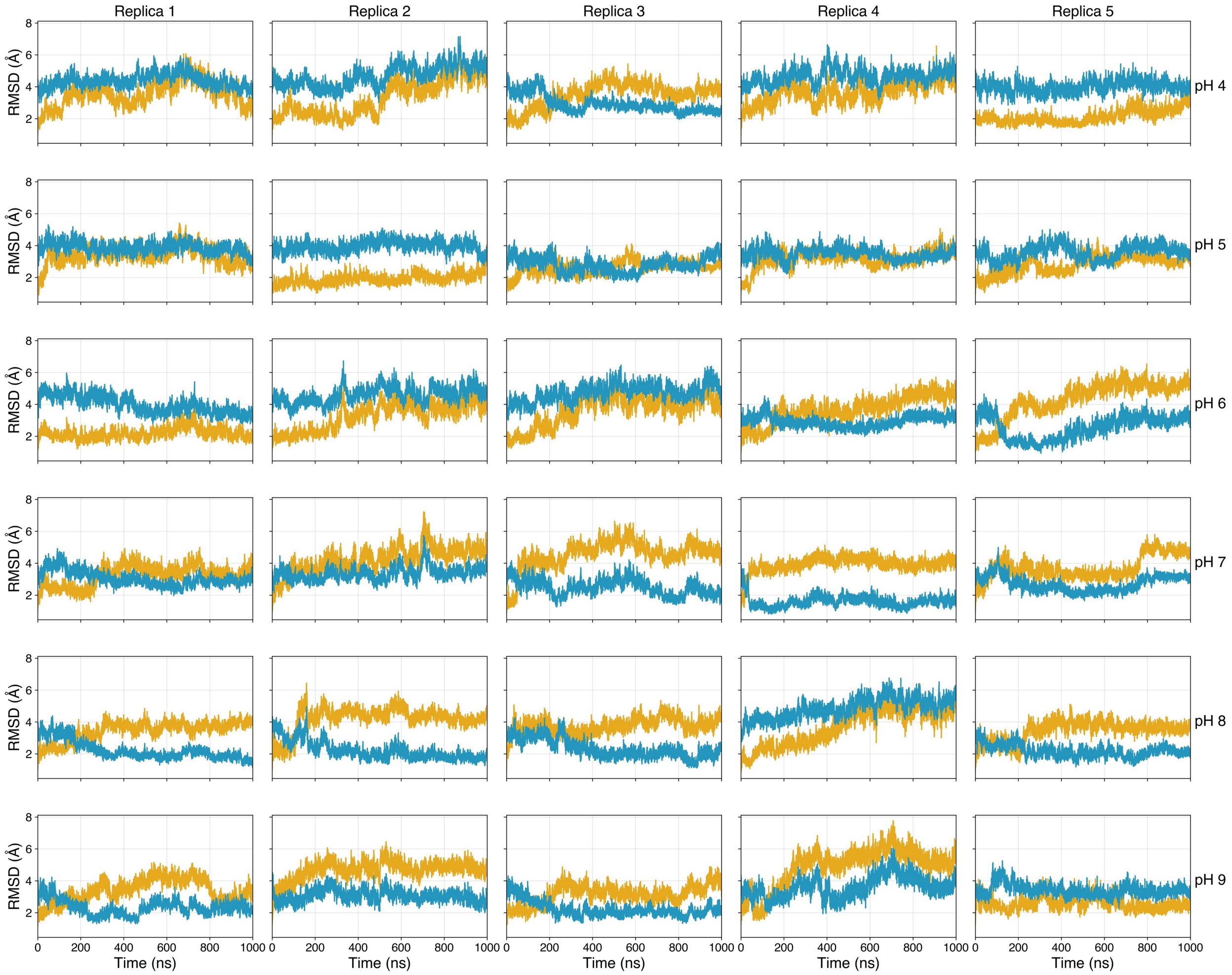


Figure S3: Root Mean Square Deviation (RMSD) of transmembrane regions across pH and replica conditions relative to active and inactive reference structures. Each subplot shows the time evolution of the RMSD values (in Å) computed for each replica (columns 1–5) at a specific pH (rows, from pH 4 at the top to pH 9 at the bottom). RMSD was calculated using GROMACS[3] over five independent molecular dynamics trajectories per pH condition. RMSD reflects the structural deviation of the receptor from two reference states: the inactive structure (PDB ID: 2RH1) shown in blue, and the active structure (PDB ID: 3P0G) shown in yellow. For each trajectory, only the C$\alpha$ atoms of TM3 (residues 102-137), TM5 (196-227), and TM6 (266-299) were used for RMSD calculation, after a least-squares alignment to the C$\alpha$ atoms of TM1 (29-61), TM2 (66-97), TM4 (146-172), and TM7 (304-328). The analysis was restricted to residues in common in both 3P0G and 2RH1 due to missing residues in the reference structures.

| pH | Replica | Ionic lock distance | Y-Y distance | NPxxY RMSD | PIF RMSD |
|---|---|---|---|---|---|
| pH 4 | replica 1 | ✗ | ✗ | ✗ | ✗ |
| | replica 2 | ✗ | ✗ | ✗ | ✗ |
| | replica 3 | ✓ | ✓ | ⚠ | ✗ |
| | replica 4 | ✗ | ✗ | ✗ | ✗ |
| | replica 5 | ✗ | ✓ | ✓ | ⚠ |
| pH 5 | replica 1 | ✗ | ✗ | ✗ | ⚠ |
| | replica 2 | ✗ | ✗ | ⚠ | ⚠ |
| | replica 3 | ✓ | ⚠ | ⚠ | ⚠ |
| | replica 4 | ✗ | ✗ | ✗ | ✗ |
| | replica 5 | ⚠ | ✗ | ✗ | ⚠ |
| pH 6 | replica 1 | ✗ | ✗ | ✗ | ✗ |
| | replica 2 | ✗ | ⚠ | ✗ | ✗ |
| | replica 3 | ✗ | ⚠ | ✗ | ✗ |
| | replica 4 | ✓ | ✓ | ✗ | ✗ |
| | replica 5 | ✓ | ✓ | ✓ | ✓ |
| pH 7 | replica 1 | ✓ | ✗ | ✗ | ⚠ |
| | replica 2 | ✓ | ✗ | ✗ | ⚠ |
| | replica 3 | ✓ | ⚠ | ✓ | ✓ |
| | replica 4 | ✓ | ✓ | ✓ | ✓ |
| | replica 5 | ✓ | ⚠ | ⚠ | ✓ |
| pH 8 | replica 1 | ✓ | ⚠ | ⚠ | ✓ |
| | replica 2 | ✓ | ⚠ | ✓ | ✓ |
| | replica 3 | ✓ | ⚠ | ✓ | ✓ |
| | replica 4 | ✗ | ⚠ | ✗ | ✗ |
| | replica 5 | ✓ | ⚠ | ✗ | ✓ |
| pH 9 | replica 1 | ✓ | ⚠ | ⚠ | ✓ |
| | replica 2 | ✓ | ⚠ | ⚠ | ✓ |
| | replica 3 | ✓ | ⚠ | ⚠ | ✓ |
| | replica 4 | ✓ | ⚠ | ⚠ | ⚠ |
| | replica 5 | ✗ | ✗ | ⚠ | ⚠ |

✗: Remains active ⚠: Partially inactivated ✓: Inactivated

Figure S4: Table of the microswitches inactivation during the performed simulations.

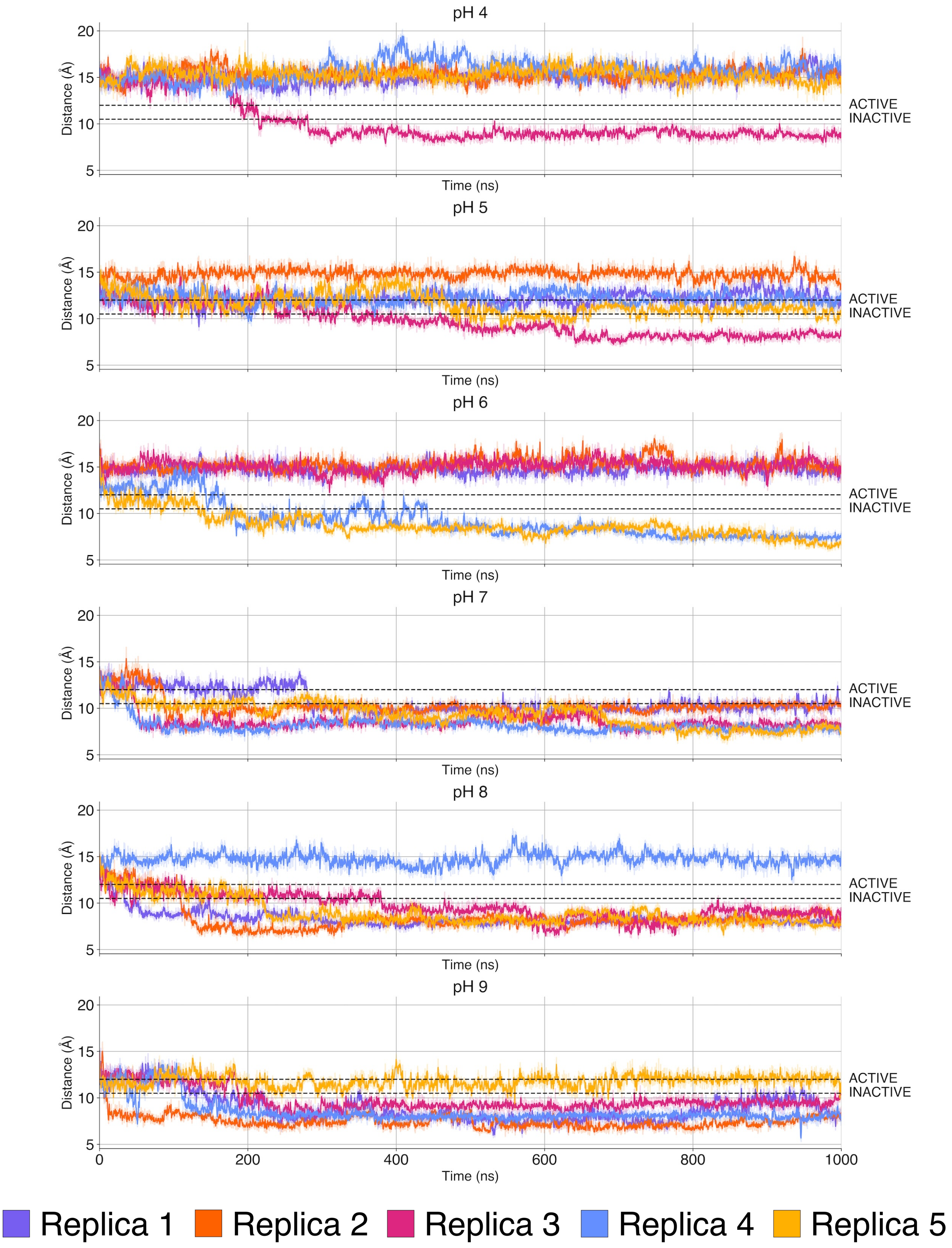


Figure S5: Time evolution for the Ionic lock distance in all the simulations performed.

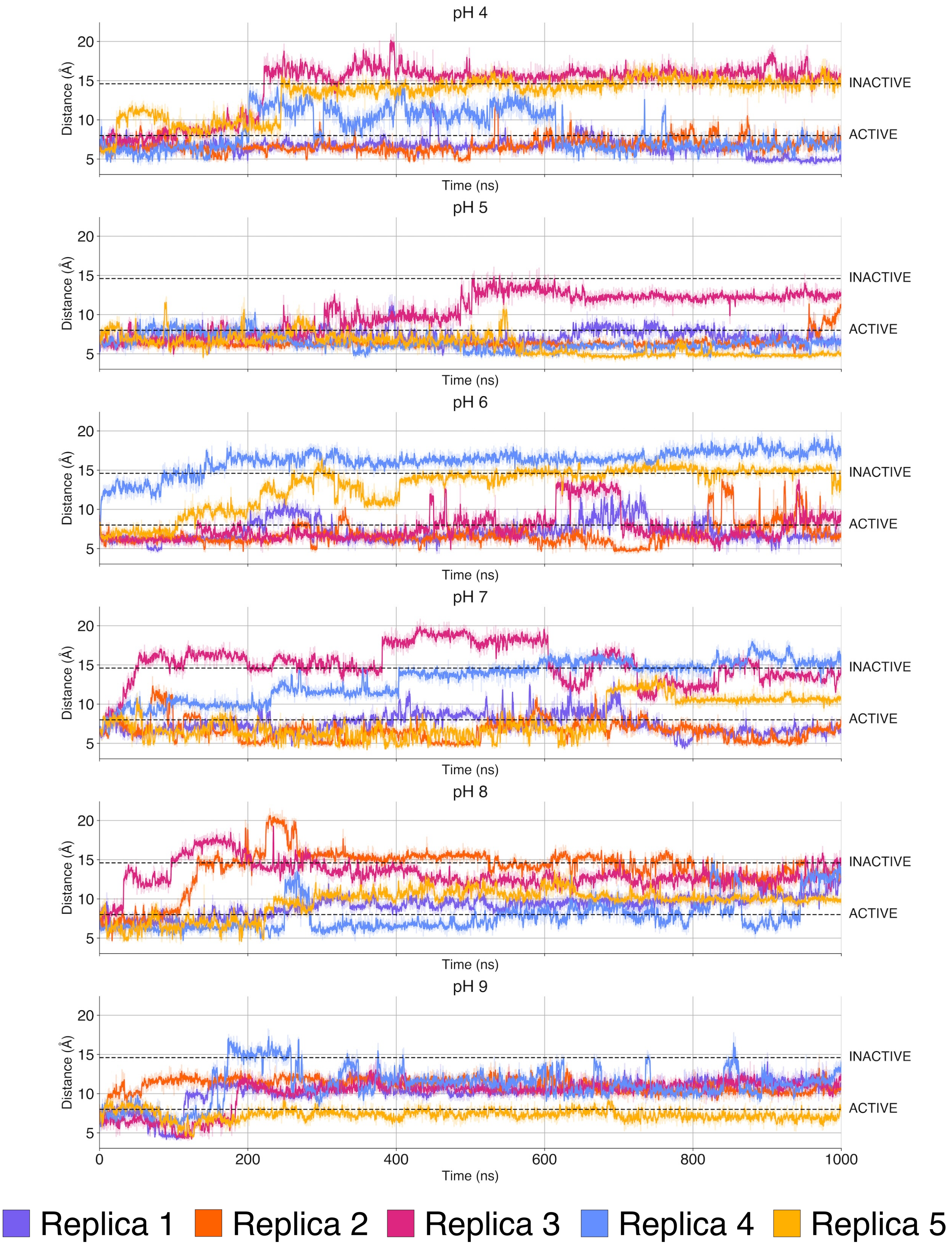


Figure S6: Time evolution for the Y-Y distance in all the simulations performed.

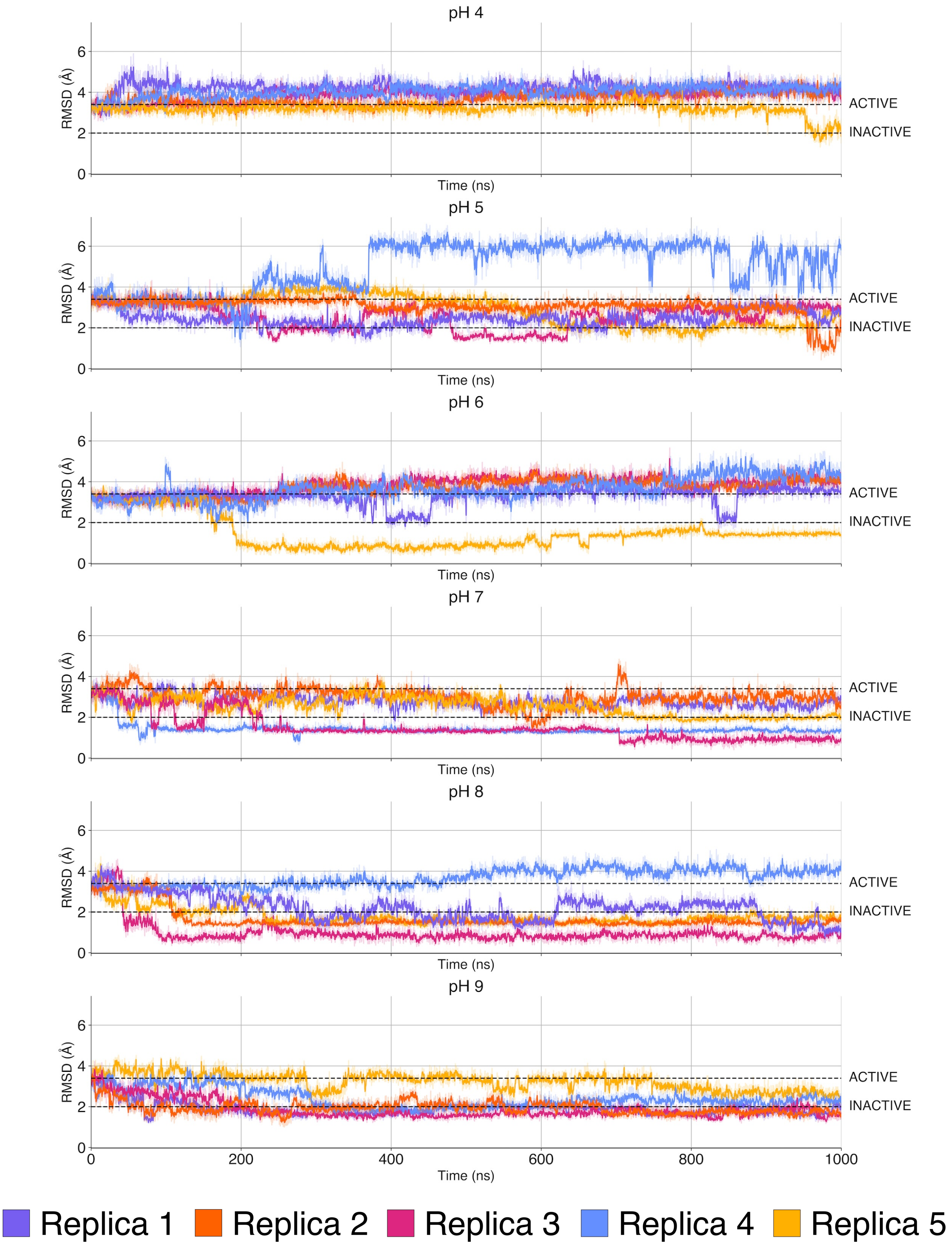


Figure S7: Time evolution for the PIF motif RMSD in all the simulations performed.

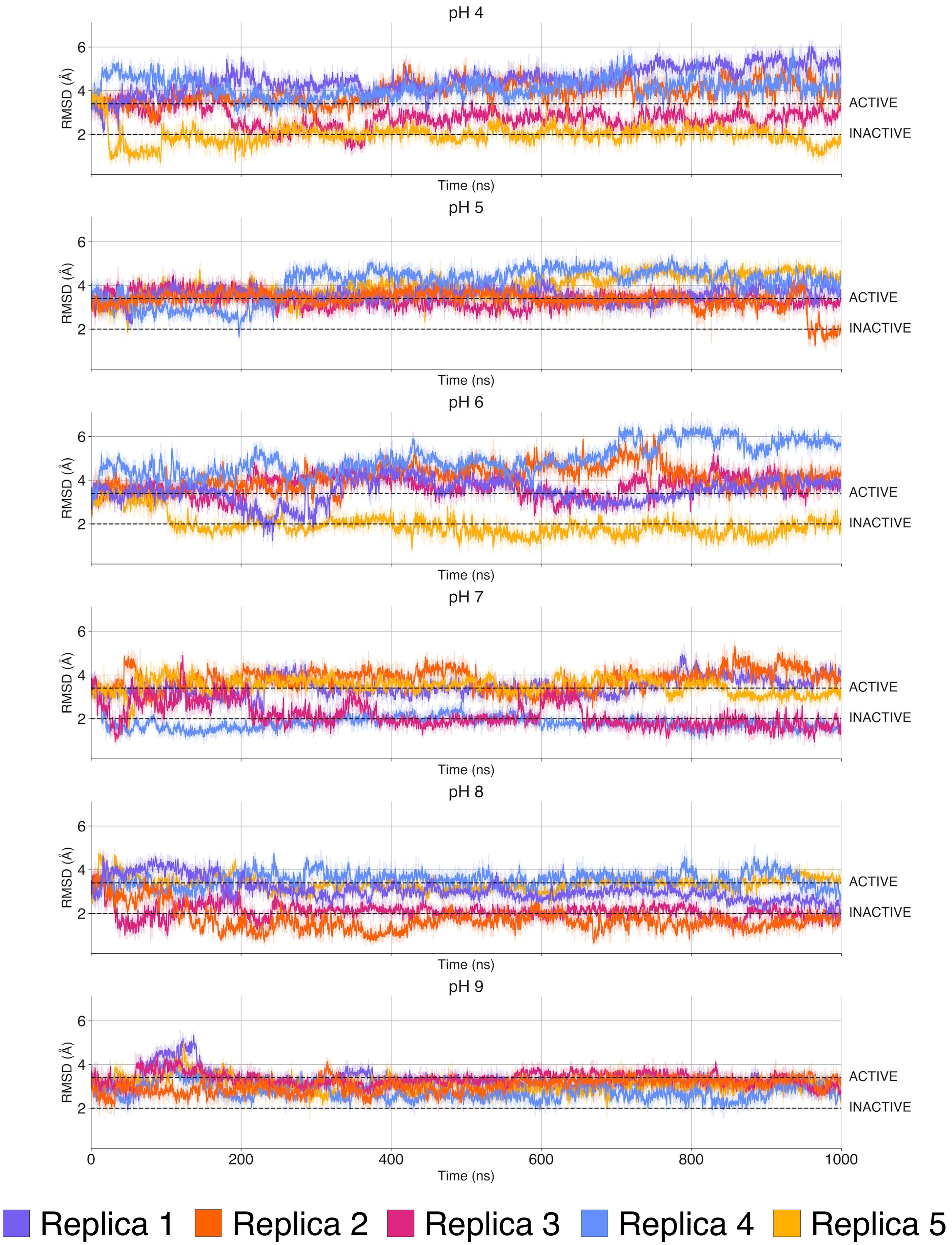


Figure S8: Time evolution for the NPxxY motif RMSD in all the simulations performed.

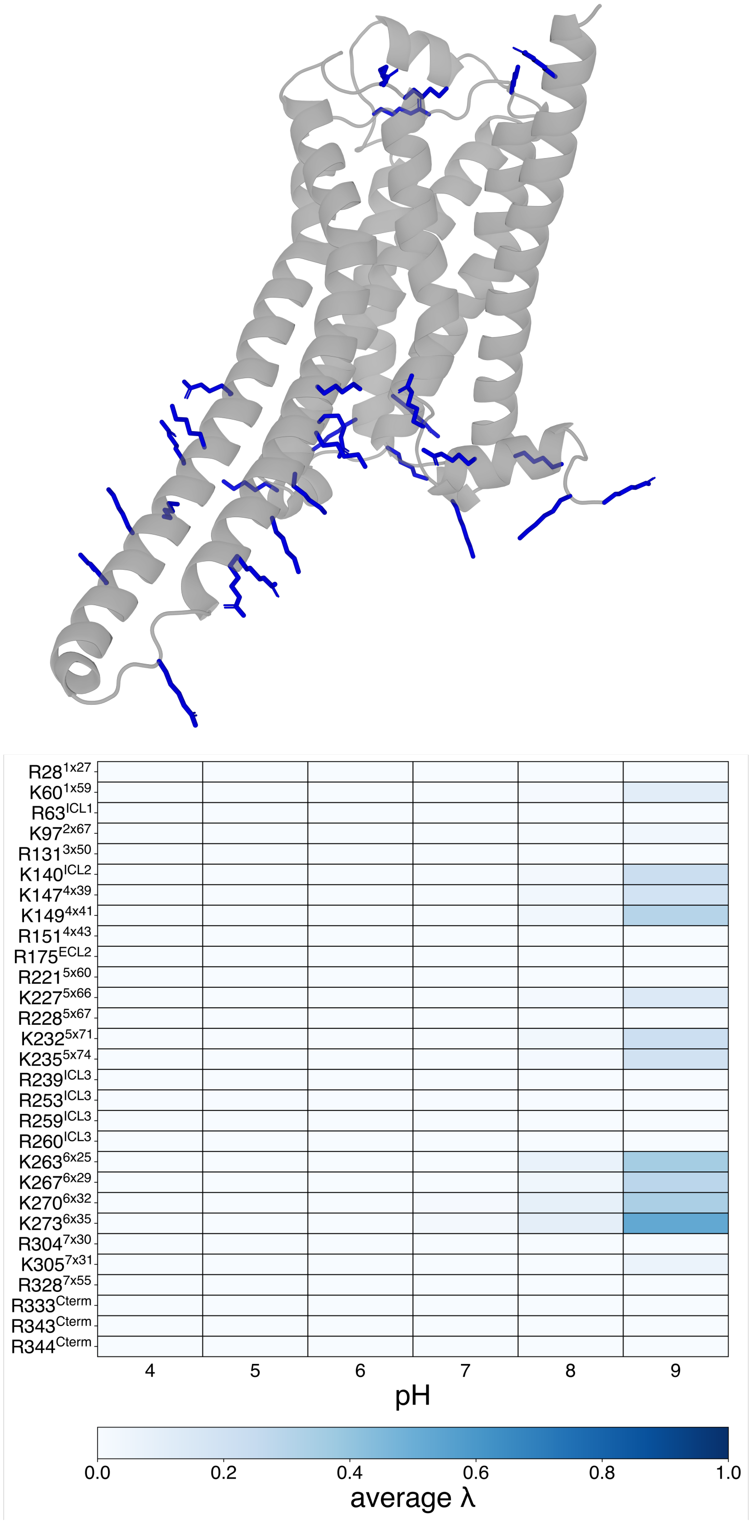


Figure S9: Titrable basic residues and their protonation state at different pH. On top, the structure of $\beta_2$AR with basic residues highlighted in blue. Below, a colormap representing the charge state of such residues. All the basic amino acids are neutral for all the simulation from pH 4 to 8. At pH 9 a small number of residues show a partial change in protonation state.

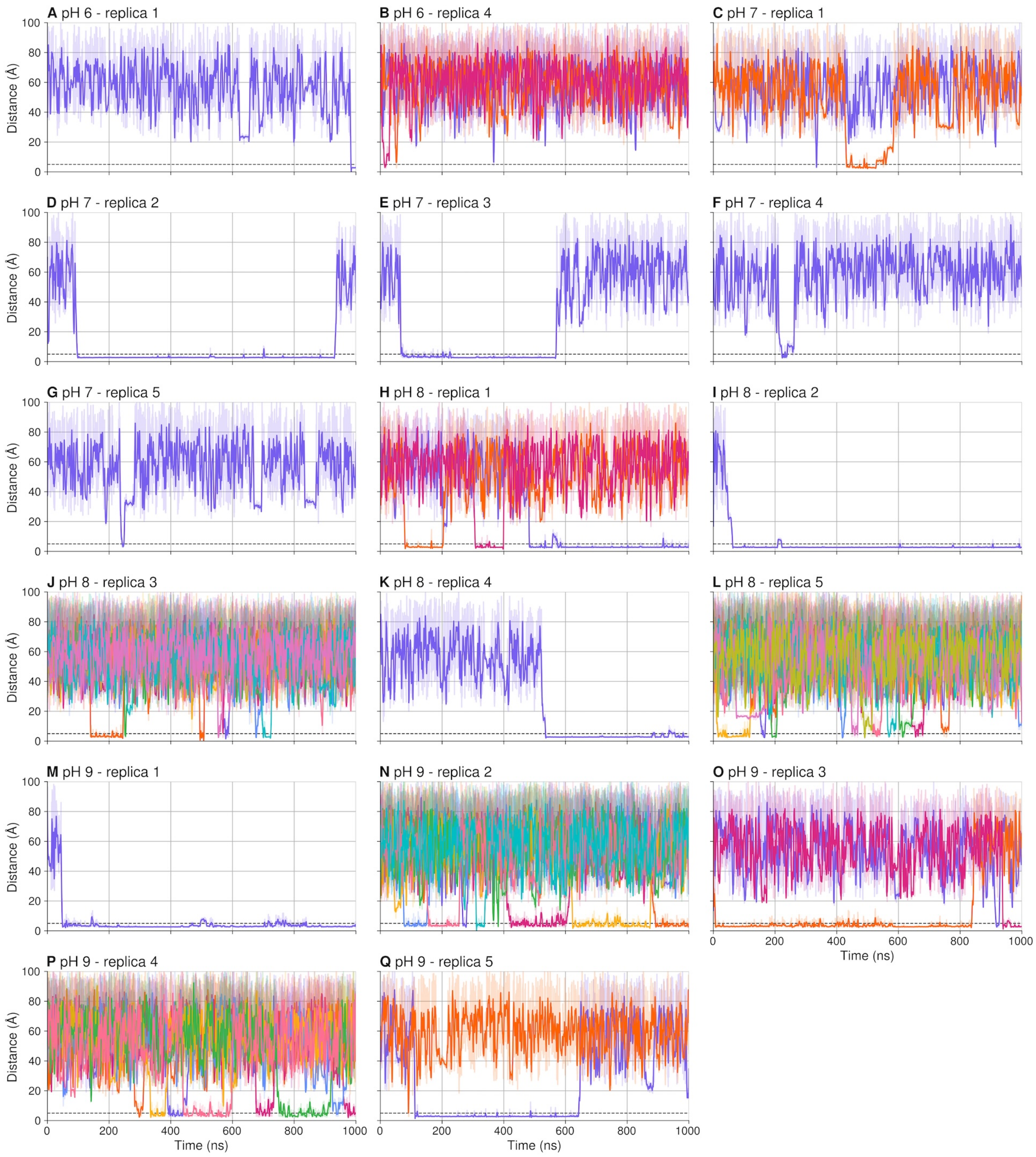


Figure S10: Time evolution of distances between $Na^+$ ions and the C$\gamma$ atom of residue D113$^{3x32}$ across different pH conditions and replicas. Each subplot corresponds to a specific pH and replica, with individual ion traces shown. Raw distance data (semi-transparent lines) are plotted alongside smoothed trajectories (solid lines) obtained using a Savitzky–Golay filter (window size = 51 frames, polynomial order = 3). The black dashed line marks the 5 Å distance threshold used to define ion-residue contact. Distances were computed from molecular dynamics trajectories using MDAnalysis,[4] and time is shown in nanoseconds. Ion traces correspond to atom numbers in the reference structure protein_na.pdb as follows:

A) 5297 in purple; B) 5303 in purple, 5323 in orange, 5330 in pink; C) 5296 in purple, 5323 in orange; D) 5330 in purple; E) 5305 in purple; F) 5294 in purple; G) 5290 in purple; H) 5279 in purple, 5291 in orange, 5331 in pink; I) 5295 in purple; J) 5281 in purple, 5300 in orange, 5303 in pink, 5304 in light blue, 5307 in yellow, 5308 in green, 5315 in coral pink, 5317 in cyan, 5332 in soft magenta; K) 5317 in purple; L) 5279 in purple, 5282 in orange, 5298 in pink, 5299 in light blue, 5306 in yellow, 5310 in green, 5316 in coral pink, 5320 in cyan, 5322 in soft magenta, 5325 in olive yellow; M) 5280 in purple; N) 5284 in purple, 5299 in orange, 5302 in pink, 5303 in light blue, 5308 in yellow, 5309 in green, 5314 in coral pink, 5317 in cyan; O) 5281 in purple, 5306 in orange, 5312 in pink; P) 5279 in purple, 5282 in orange, 5294 in pink, 5307 in light blue, 5321 in yellow, 5324 in green, 5327 in coral pink; Q) 5316 in purple, 5332 in orange.

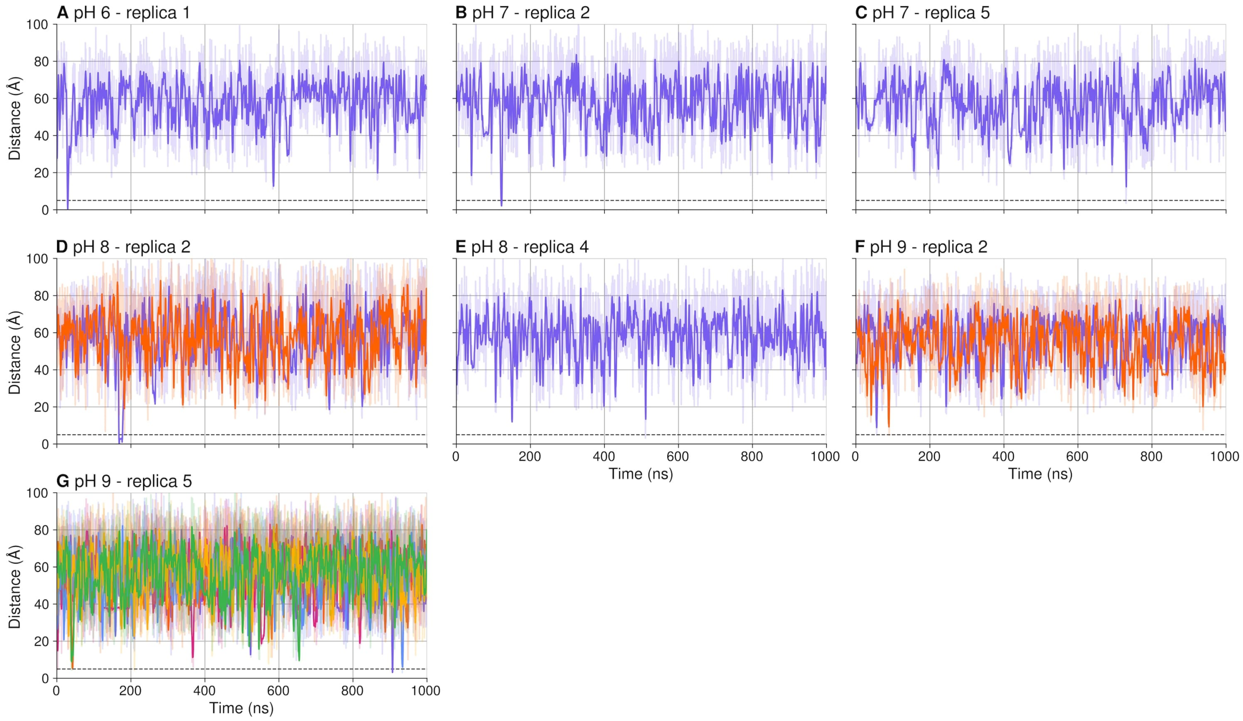


Figure S11: Time evolution of distances between $Na^+$ ions and the CG atom of residue $D130^{3x49}$ across different pH conditions and replicas. Each subplot corresponds to a specific pH and replica, with individual ion traces shown. Raw distance data (semi-transparent lines) are plotted alongside smoothed trajectories (solid lines) obtained using a Savitzky–Golay filter (window size = 51 frames, polynomial order = 3). The black dashed line marks the 5 Å distance threshold used to define ion-residue contact. Distances were computed from molecular dynamics trajectories using MDAnalysis,[4] and time is shown in nanoseconds. Ion traces correspond to atom numbers in the reference structure protein_na.pdb as follows: A) 5315 in purple; B) 5304 in purple; C) 5298 in purple; D) 5306 in purple, 5307 in orange; E) 5288 in purple; F) 5289 in purple, 5326 in orange; G) 5293 in purple, 5314 in orange, 5315 in pink, 5317 in light blue, 5319 in yellow, 5329 in green.

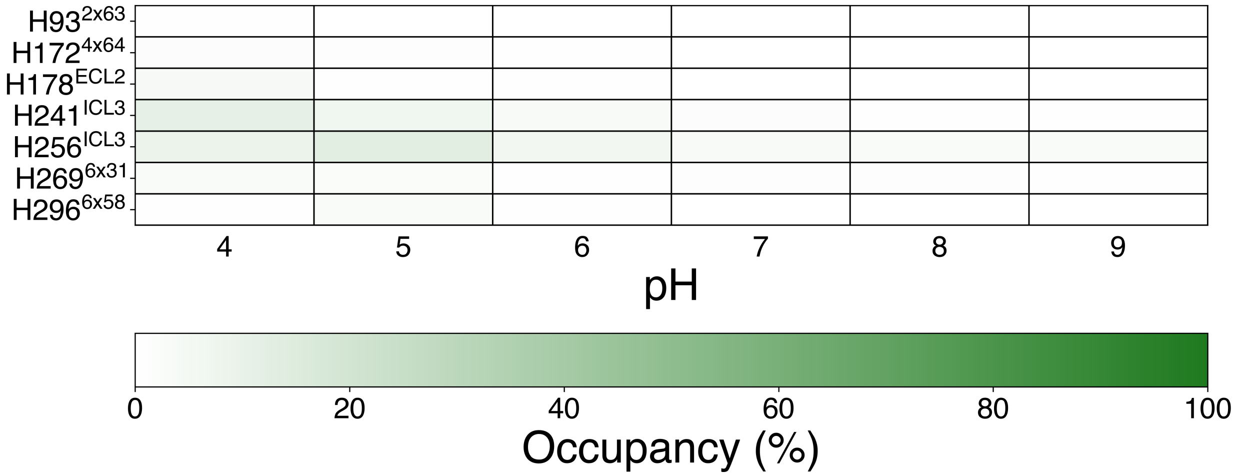


Figure S12: Occupancy of $Cl^-$ ions to histidine residues. In all the pH conditions studied here we do not see a lasting binding between chlorine and charged histidine residue: the only cases where the time percentage overcomes 10% is for pH between 4 and 5, with residues in the ICL3 completely exposed to solvent (H241$^{ICL3}$ and H256$^{ICL3)}$.
